# A Germinal Center-Associated Microenvironmental Signature Reflects Malignant Phenotype and Outcome of Diffuse Large B-cell Lymphoma

**DOI:** 10.1101/833947

**Authors:** Kohta Miyawaki, Koji Kato, Takeshi Sugio, Kensuke Sasaki, Hiroaki Miyoshi, Yuichiro Semba, Yoshikane Kikushige, Yasuo Mori, Yuya Kunisaki, Hiromi Iwasaki, Toshihiro Miyamoto, Frank C. Kuo, Jon C. Aster, Koichi Ohshima, Takahiro Maeda, Koichi Akashi

## Abstract

Diffuse large B-cell lymphoma (DLBCL) is the most common B-cell malignancy with varying prognosis after the gold standard rituximab, cyclophosphamide, doxorubicin, vincristine, and prednisone (R-CHOP). Several prognostic models have been established by focusing primarily on characteristics of lymphoma cells themselves including cell-of-origin, genomic alterations, and gene/protein expressions. However, the prognostic impact of lymphoma microenvironment and its association with characteristics of lymphoma cells are not fully understood. Using highly-sensitive transcriptome profiling of untreated DLBCL tissues, we here assess the clinical impact of lymphoma microenvironment on the clinical outcomes and pathophysiological, molecular signatures in DLBCL. The presence of normal germinal center (GC)-microenvironmental cells, including follicular T cells, macrophage/dendritic cells, and stromal cells, in lymphoma tissue indicates a positive therapeutic response. Our prognostic model, based on quantitation of transcripts from distinct GC-microenvironmental cell markers, clearly identified patients with graded prognosis independently of existing prognostic models. We observed increased incidences of genomic alterations and aberrant gene expression associated with poor prognosis in DLBCL tissues lacking GC-microenvironmental cells relative to those containing these cells. These data suggest that the loss of GC-associated microenvironmental signature dictates clinical outcomes of DLBCL patients reflecting the accumulation of “unfavorable” molecular signatures.

## Introduction

Diffuse large B-cell lymphoma (DLBCL) is the most common subtype of B-cell non-Hodgkin lymphomas with heterogeneous clinicopathologic features. By performing global gene expression profiling (GEP), Alizadeh et al. have grouped DLBCL cases into two subtypes based on the cell-of-origin (COO) of lymphoma cells. The germinal center B-cell-like (GCB) type exhibits the signature of B-cells in germinal center (GC) of normal secondary lymphoid organs, while lymphoma cells of the activated B-cell (ABC)-like type resemble post-GC B-cells that transit from the GC for plasmacytic differentiation (1). Among DLBCL patients treated with multi-agent chemotherapy consisting of cyclophosphamide, doxorubicin, vincristine, and prednisone (CHOP), patients with ABC type disease generally show significantly worse prognosis than do those with GCB type (1).

Other groups have proposed immunohistochemistry (IHC)-based algorithms for formalin-fixed paraffin-embedded (FFPE) tissues that recapitulate microarray-based COO classification (e.g. Hans’ criteria) (2, 3). In clinical trials of patients undergoing R-CHOP (CHOP plus Rituximab, a chimeric antibody against the CD20 B-cell antigen) regimens, IHC-based COO criteria have minimal prognostic power (4–8). Given that R-CHOP still remains the gold standard after numerous attempts for improving R-CHOP such as intensification of rituximab or CHOP, introduction of next-generation anti-CD20 antibodies, and addition of novel therapeutic agents (e.g. proteasome inhibitor) (9), the accurate stratification model is still in demand which could predict clinical outcomes after R-CHOP. A recent large-scale genomic study of >1,000 DLBCL patients revealed the mutational heterogeneity of DLBCL and concluded that the best performing predictive model could be only be achieved by combining DNA- and RNA-risk models (10). Currently, the International Prognostic Index (IPI) remains the most reliable prognostic model, although it is based solely on clinical variables and patient status. Thus, a more accurate prognostic model, one that comprehensively reflects DLBCL pathophysiology and helps physicians in therapeutic decision making, is still needed.

As naïve B-cells develop into antibody-secreting plasma cells in lymph nodes (LNs), they undergo stage-specific genome editing activities to generate variable immunoglobulins. Although required for immune cell diversity, this process predisposes B-cells to transformation if the editing machinery is hijacked (11). Dynamic processes of B-cell development in the GC are controlled by cell-intrinsic activities and by surrounding microenvironmental cells, which include follicular T cells, stromal cells, dendritic cells (DCs) and macrophages. Microenvironmental cells are in fact necessary for GC formation and are recruited from the periphery in response to normal immune signals that govern GC formation (12, 13). These observations suggest that the tumor microenvironment may play a role in DLBCL pathogenesis.

## Results

### Transcriptome profiling of untreated DLBCL tissues identifies prognostic genes

We collected 280 FFPE tissues from de novo DLBCL patients (30, 170 and 80 cases for pilot, training and validation cohort, respectively) (Fig. 1A). Cases of relapsed or transformed disease were excluded. All the patients had been treated with R-CHOP or R-THPCOP (tetrahydropyranyladriamycin, cyclophosphamide, vincristine, and prednisone) (14). Median observation time for surviving patients was 3.67 and 5.60 years, and 3-year disease-free survival (DFS) rate was 66.3% and 65.0% in the training and validation cohorts, respectively. Additional patient characteristics are provided in table S1. To quantify fragmented mRNA in FFPE tissues, we employed the nCounter system, which enables highly sensitive and accurate RNA detection (15–17). In brief, we compared two replicated measurements of 48 representative genes (table S2) with each other in an FFPE sample. The result showed exceptionally high measurement reproducibility, suggesting that this screen could detect all RNAs expressed in specimens whether they are from lymphoma cells or rare microenvironment components (fig. S1A).

**Fig. 1.**
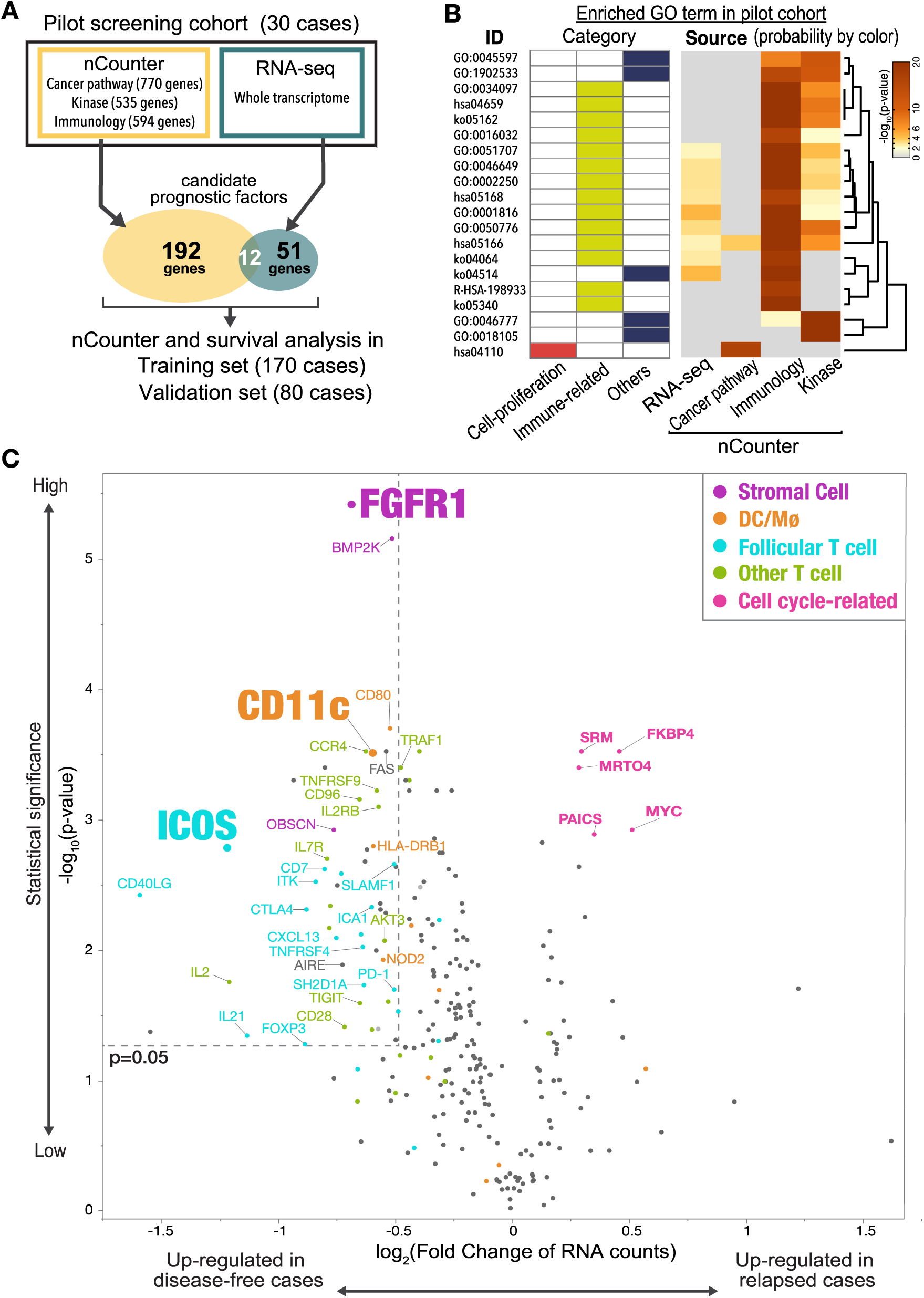
Identification of prognostic factors for DLBCL outcomes. (**A)** Schematic of overall study design. Thirty cases of newly-diagnosed DLBCL were recruited for a pilot screen using nCounter and RNA-seq methods, followed by analysis of two larger cohorts. (**B**) Gene Ontology (GO) analysis of candidate prognostic genes from the pilot screen, which was performed using the Metascape web application (http://metascape.org). Enriched GO terms are shown by ID, category, and derived pilot cohort. Probabilities are depicted by color. Most prognostic genes were linked to immune-related terms, irrespective of source. (**C)** A volcano plot indicates differentially expressed genes as favorable (left) and poor (right) prognostic factors in the training cohort. Most unfavorable indicators were cell cycle-related (magenta). Note that many microenvironmental cell-related genes were associated with favorable prognosis. A Mann-Whitney U test (unpaired) with bootstrap was performed to calculate P-values.

We performed a pilot screen to identify genes correlated with specific patient outcomes by analyzing FFPE samples obtained at initial diagnosis from 30 DLBCL patients: 15 showing favorable outcomes (>4 years in remission) and 15 showing poor prognosis (primary refractory or early relapse disease). We used commercially available probe sets that included 1899 genes related to immunology, cancer, or kinase pathway panels (Fig. 1A). This screen identified 204 genes with a statistically significant association with either favorable or poor prognosis (q-value < 0.05, fig. S1B and tables S3-5). More than half of candidate prognostic genes was derived from the Immunology panel, including many T cell-related genes, such as those related to follicular T cells (fig. S1B). To cover the limitation in the number of genes analyzed, we performed whole transcriptome analysis on the 30 samples of the pilot screen using RNA sequencing (RNA-seq) and extracted an additional 51 differentially-expressed genes as candidate prognostic markers (table S6). Gene ontology (GO) analysis revealed that most of prognostic genes were associated with immune responses, while only one cell proliferation-related term was enriched, suggesting that immune signature has a significant effect on DLBCL clinical outcomes (Fig. 1B). Transcript levels of each of the twelve genes (*ICOS*, *CD80*, *CTLA4*, *EGR2*, *CD58*, *TRAF1*, *C1QBP*, *IL21*, *CD40LG*, *FAS*, *MYC*, *CD96*) were identified as prognostic factors both in nCounter and RNA-seq analyses. We added additional 192 genes for reference and normalization: COO-defining genes, reported prognostic genes, B-cell-associated genes, and microenvironmental cells-specific genes, and designed a custom probe set containing a total of 447 genes for nCounter analysis (table S7).

We next applied similar analysis to 170 DLBCL samples from patients who had undergone R-CHOP-based regimens. Among these, tissue specimens came from LNs of 100 patients and extranodal sites of 70 others (fig. S1C). The scatter plot shown in Fig. 1C depicts genes upregulated in specimens representing poor (relapsed) versus favorable (disease-free) outcomes. Among genes up-regulated in patients with poor prognosis (genes depicted in the right upper area in Fig. 1C) were those related to cell proliferation, including *MYC* and its targets: *FKBP4* (18, 19), *SRM* (19), and *PAICS* (20). All four of these genes were expressed at high levels in DLBCL cell lines relative to normal LN samples (fig. S2A). As expected, MYC protein levels were proportional to *MYC* transcript levels and correlated with levels of proliferation-associated genes (fig. S2B). Kaplan-Meier curves of DFS confirmed that high expression of each proliferation-associated gene marks patients with poor prognosis (fig. S2C).

### Immune microenvironment-related genes define favorable prognosis in DLBCL

Most genes in the analysis of 170 DLBCL samples described above were identified as favorable prognostic factors (Fig. 1C, upper left box: p < 0.05 and log_2_[fold change] < −0.5). These factors were again linked to immune-related GO terms, especially those related to T-B interactions (fig. S3A). Of note, most favorable prognostic factors mark GC-related microenvironmental cells. Among them were *ICOS*, *CXCL13*, *CD40LG*, *PD-1*, *SH2D1A (SAP)*, *IL2RB*, *IL21* and *SLAMF1*, all of which are representative markers for CD4^+^ follicular T cells and regulate B-cell development in the normal GC (12, 13, 21, 22). In addition, *CD11c* and fibroblast growth factor 1 (*FGFR1*) are expressed primarily in DC/macrophage and interstitial fibroblasts/stromal cells, respectively. None of above genes were expressed in DLBCL cell lines (Fig. 2A), strongly suggesting they are derived from non-lymphoma cells. Patients with DLBCL whose specimens show high expression of these microenvironmental cell-related transcripts exhibit a significantly (*P* < 0.005) more favorable prognosis (Fig. 2B).

**Fig. 2.**
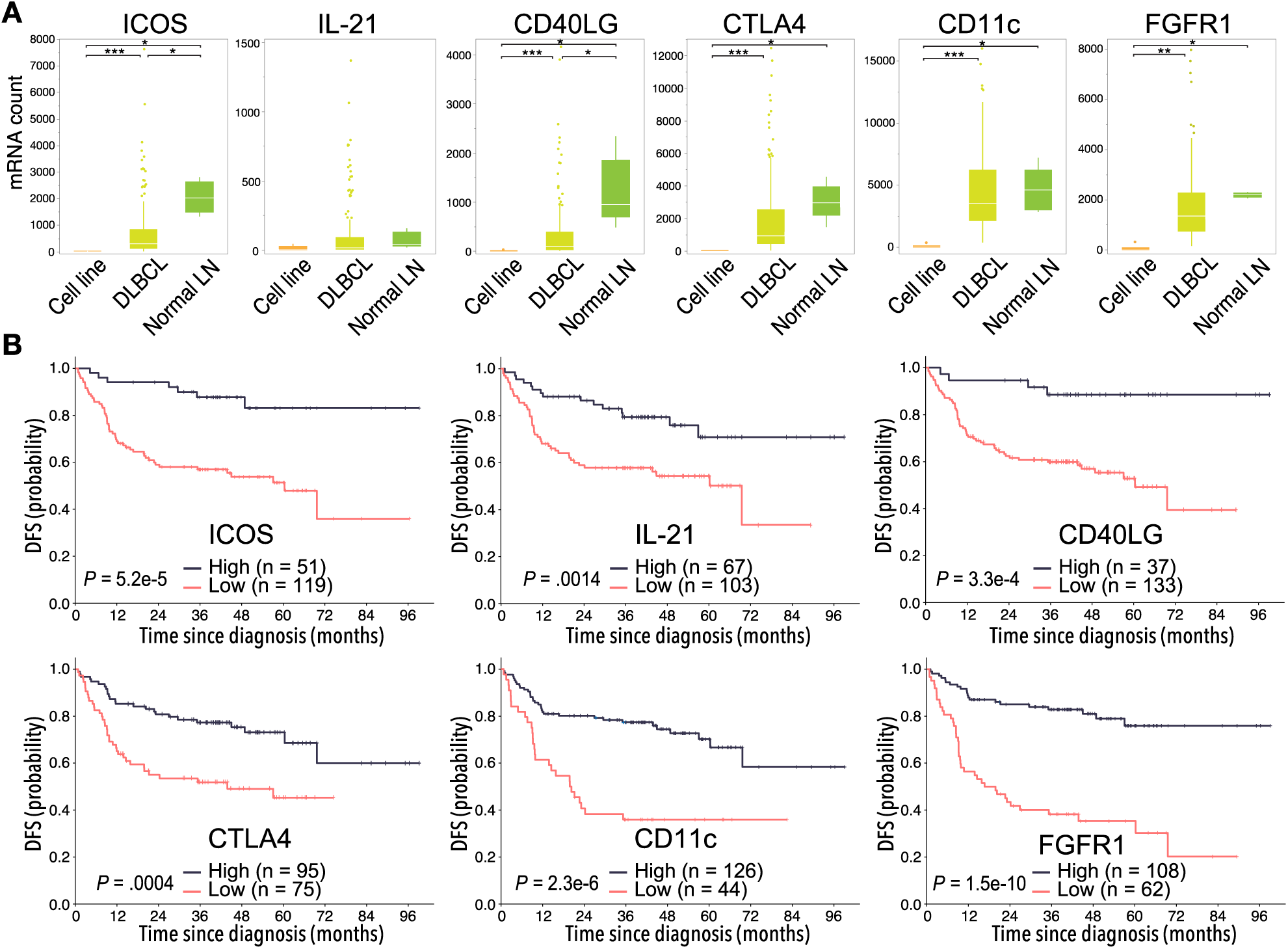
Clinical impact of extracted favorable prognostic genes. (**A)** Expression of 6 favorable prognostic genes in DLBCL cell lines (n=6), primary DLBCL specimens (n=170), and normal lymph nodes (LNs) (n=5), as shown in a Box and Whisker plot (dots show outliers). Cell lines analyzed were RC-K8, SU-DHL-1, SU-DHL-4, SU-DHL-9, Karpas-422, and OCI-Ly-3. Lymphoma lines showed no or little candidate gene expression, suggesting genes are expressed in microenvironmental cells. Significance was determined using the Steel-Dwass test (*p<0.05, **p<0.005, and ***p<0.0005). (**B**) Kaplan-Meier curves show duration of DFS based on expression levels of favorable candidate genes. P-values were calculated by log-rank test.

### Favorable prognostic genes derive from specific lymphoma microenvironment cells

To characterize T cells infiltrating lymphoma tissues, we performed single-cell GEP of CD4^+^ T cells in six cryo-preserved DLBCL specimens (Fig. 3A). More than three fourth of single CD4^+^ T cells (76.6 ± 9.7%, n=6) from favorable cases expressed the follicular T cell marker *ICOS* (Fig. 3, B and C).

**Fig. 3.**
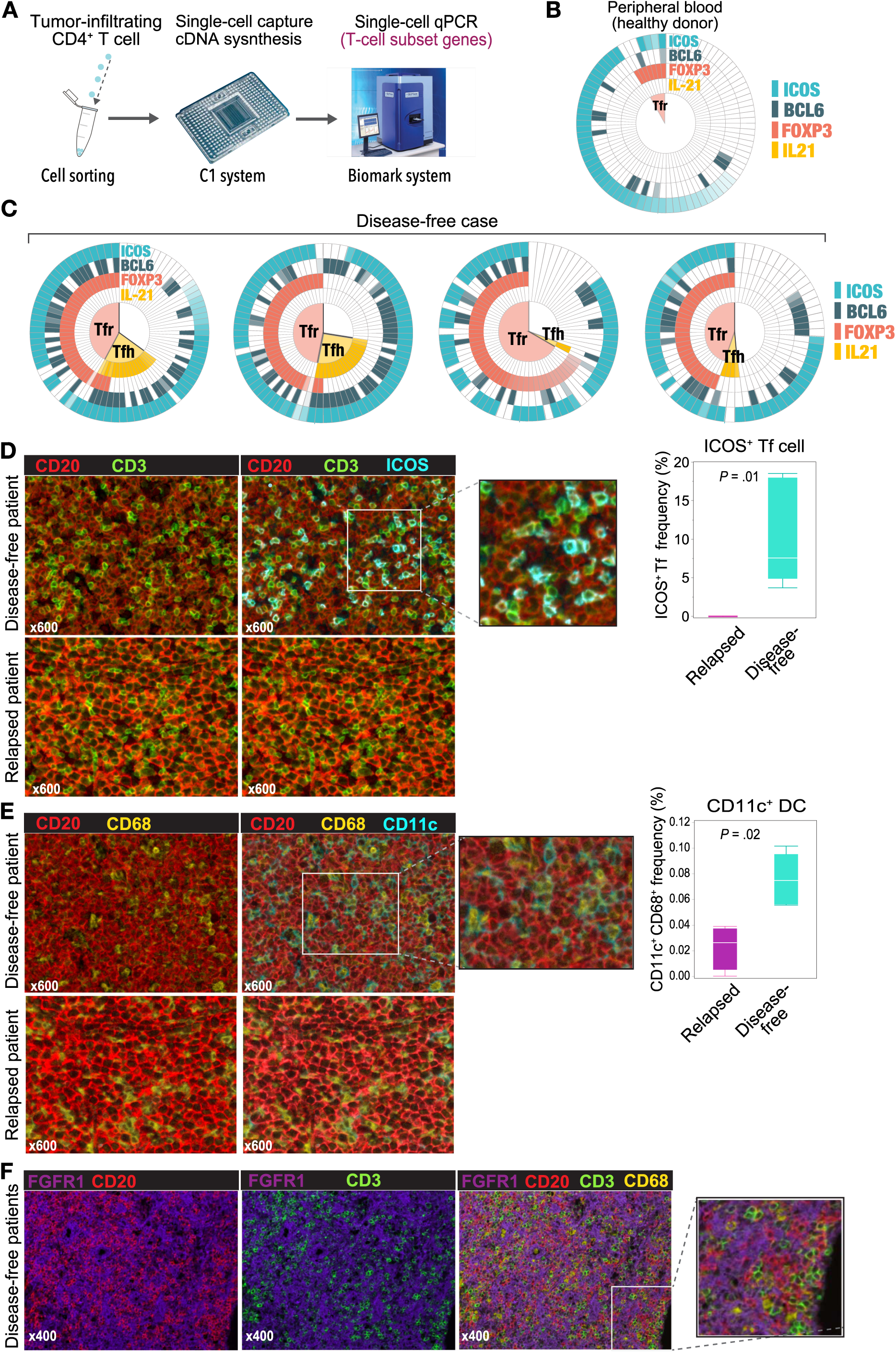
Identification of microenvironmental cell subtypes by single-cell expression profiling and imaging analysis. (**A)** Schematic representation of the experimental procedure. (**B, C**) Multi-level pie-charts show results of single-cell gene expression profiling of CD4^+^ T cells sorted from DLBCL tissues from patients with favorable prognosis (**C**) and from peripheral blood of healthy donors (controls) (**B**). Color intensity shows gene expression levels. Shown are four representative cases of six analyzed. (**D**) Left, detection of follicular T (Tf) cells in DLBCL tissue based on indicated markers using multiplexed immunofluorescence imaging analysis. Representative images taken from 10 patient samples analyzed are shown. Right, Box and Whisker plot (dots indicate outliers) shows frequency of ICOS^+^ Tf cells, which were digitally counted in five fields. P values were calculated using the Wilcoxon Rank Sum Test. (**E**) Left, comparable imaging identified CD68^+^ dendritic cells (DC)/macrophages (yellow) expressed CD11c (blue) in favorable prognostic case. Right, the proportion of CD11c^+^ CD68^+^ cells among total cells in 5 fields, shown as a Box and Whisker plot (dots indicate outliers), in relapsed versus disease-free cases, as calculated by the Wilcoxon Rank Sum Test. (**F**) Characterization of FGFR1+ cells (purple) based on staining with the lineage-specific markers CD3 (T cells), CD68 (DC/macrophages), and CD20 (B-cells).

About half of *ICOS*^+^ CD4^+^ T cells co-expressed *BCL6*, follicular helper T (Tfh) cell master transcription factor (23, 24), as well as *PD-1*, *SAP* and *OX40* (25). Interestingly, some of Tfh gene-expressing cells expressed *FOXP3* in a manner mutually exclusive of the Tfh cytokine *IL21*, suggesting that those cells were follicular regulatory T (Tfr) cells (Fig. 3C).

We next used multispectral fluorescence imaging to identify the origin of favorable prognostic genes in the DLBCL tissues. Infiltrating CD3^+^ ICOS^+^ follicular T cells (double-stained by cyan and green in Fig. 3D, left) scattered around CD20^+^ lymphoma cells (red) in DLBCL tissues. Most CD3^+^ T cells (green) express ICOS protein (cyan) in disease-free cases, while not in relapsed cases. Note that the frequencies of CD20^+^ lymphoma B-cells and overall CD3^+^ T cells were comparable between them. The number of follicular T cells in tissues from disease-free patients exceeded that in tissues from relapsed patients (*P* = 0.01, Fig. 3D, right). Most CD11c-positive cells co-expressed pan-macrophage marker CD68, but not CD3 or CD20, and exhibited a “dendritic” morphology (Fig. 3E). CD11c^+^ CD68^+^ DC/macrophages were also enriched in samples from disease-free patients (*P* = 0.02), while CD68^+^ DC/macrophages did not express CD11c in relapsed case. In contrast, the expression of FGFR1, which is usually expressed by interstitial fibroblast or stromal cells (26) did not merge with any lineage-specific markers such as CD3, CD11c, CD20 or CD68, suggesting that they are tertiary microenvironment components (Fig. 3F). We also performed multispectral imaging in normal second lymphoid tissues to assess ICOS, CD11c, and FGFR1 protein levels (fig S3, B and C). Interestingly, all three proteins were expressed reactive LNs, preferentially in the GC region, but not in steady-state B-cell follicles. Overall, these observations support the idea that the presence in DLBCL specimens of microenvironmental cells seen in the normal GC is associated with favorable clinical outcomes (12, 27).

### Immune microenvironment signature dictates DLBCL prognosis independently of existing stratification models

*ICOS*, *CD11c*, or *FGFR1* mRNAs were the top-ranked predictors of favorable prognosis with the smallest P-values in log-rank test among three different microenvironmental cell components (Fig. 1C and 2B). Thus, we asked whether these markers comprised a GC-associated microenvironmental signature that could predict patient outcomes. To evaluate this, we developed a scoring system that we call a DLBCL Microenvironmental Signature (DMS) based on *ICOS*, *CD11c*, and *FGFR1* expression (Fig. 4A). We then constructed Kaplan-Meier curves for DFS of patients segregated by DMS score (Fig. 4B). Patients with DMS scores of 3, 2, 1 and 0 points exhibited statistically significant differences in probability of 3-year DFS survival of 0.903 (*P* = 1.2e-10, 95% confidence interval [CI], 0.818-0.998), 0.807 (95% CI, 0.710-0.917), 0.514 (95% CI, 0.375-0.703) and 0.246 (95% CI, 0.131-0.462), respectively. These significant differences were evident even among patients with extranodal DLBCL lesions (*P* = 2.1e-5, Fig. 4C), cases in which microenvironmental cells must migrate to lymphoma tissue. The DMS score also predicted overall survival (OS) in nodal and extranodal cases (*P* = 2.0e-6, fig. S4A). Of note, expression levels of B cell-related genes were comparable regardless of DMS score, suggesting no difference in terms of tumor content across samples (fig. S4B). Patients of GCB or non-GCB types based on the Hans COO classification were distributed equally among different DMS scores (*P* = 0.6627, Fig. 4D). In contrast, based on Lymph2Cx analysis, which is a refined COO model based on the nCounter assay (28), GCB and ABC types more frequently corresponded to DMS-high and DMS-low cases, respectively (*P* = 3.0e-4, Fig. 4E). Importantly, the DMS score could stratify patients with different prognoses within each COO type (Fig. 4, D and E), suggesting that the DMS score is a powerful prognostic indicator independent of COO. Patients with high or low IPI also distributed equally into groups with different DMS scores (*P* = 0.0548). In both IPI low and high groups, higher DMS scores clearly marked patients with better outcomes (*P* = 0.0065 and 0.00021, Fig. 4F).

**Fig. 4.**
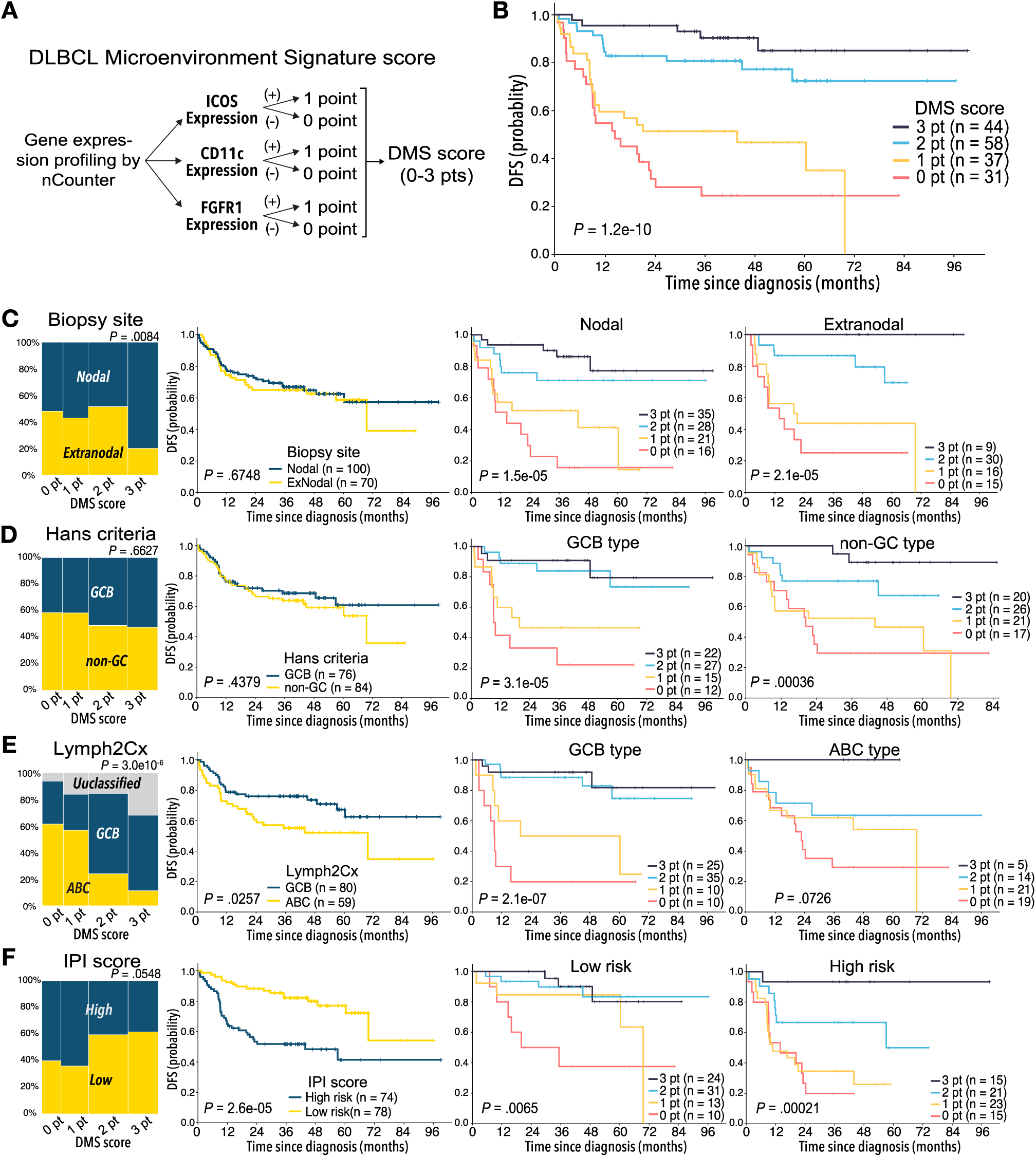
Clinical impact of DLBCL microenvironment-based stratification model. (**A)** Schematic showing algorithm used to calculate DLBCL microenvironment signature (DMS) score. (**B**) Kaplan-Meier DFS curve based on DMS score. (**C-F**) Mosaic plots showing correlation of DMS score with disease site and canonical prognostic models: (**C**) disease site (n=170), (**D**) Hans criteria (n=160), (**E**) Lymph2Cx (n=170), (**F**) IPI score (n=152). The correlation was calculated using the Fisher’s exact test. DMS-low cases were enriched in ABC type DLBCL, based on Lymph2Cx, and vice versa, but we observed no correlation with Hans criteria or IPI score. Shown are Kaplan-Meier DFS curves based on disease site, Hans criteria, Lymph2Cx (31 unclassified cases were excluded), and IPI in all cases and by DMS score in each subgroup. The DMS score had prognostic value in all subtypes, based on these classifications. A log-rank test was used for survival analysis.

### The prognostic impact of DMS is validated by independent cohorts

We next validated the prognostic value of the DMS score using an independent cohort of 80 de novo DLBCL cases (Fig. 1A and table S1). As expected, the DMS score defined by the same cut-off values as training cohort successfully delineated clinical outcomes of patients receiving R-CHOP therapy: patients with different DMS scores exhibited statistically significant differences in probability of DFS (*P* = 2.1e-5, Fig. 5A) and OS (*P* = 2.1e-5, Fig. 5B). Multivariate analysis revealed that the DMS score is a prognostic indicator independent of IPI criteria and Lymph2Cx (*P* = 0.0283, table S8). We also validated the DMS score using publicly-available RNA-seq datasets of two large cohorts, composed of 229 and 604 DLBCL cases (10, 29). Patients with DLBCL whose specimens showed high expression of DMS-related transcripts exhibited a significantly better prognosis (*P* < 0.005, fig. S5A), and the DMS score further stratified patients with different prognosis in these cohorts (*P* = 2.1e-11 and 3.5e-7, fig. S5B). The DMS score appeared to be independent of prognostic factors identified by these studies: there was a trend toward inferior OS among patients with low DMS score as compared with those with high score in each disease subgroup (fig. S6). Furthermore, multivariate analysis revealed independence of DMS score from these prognostic factors (*P* = 2.18e-4 and 1.26e-4, table S9 and S10). Of note, the DMS score stratified the “genetically unclassifiable group”, which consists of more than 50% of reported cases (29) into clinically-distinct subgroups (*P* = 0.0026, fig. S6A).

**Fig. 5.**
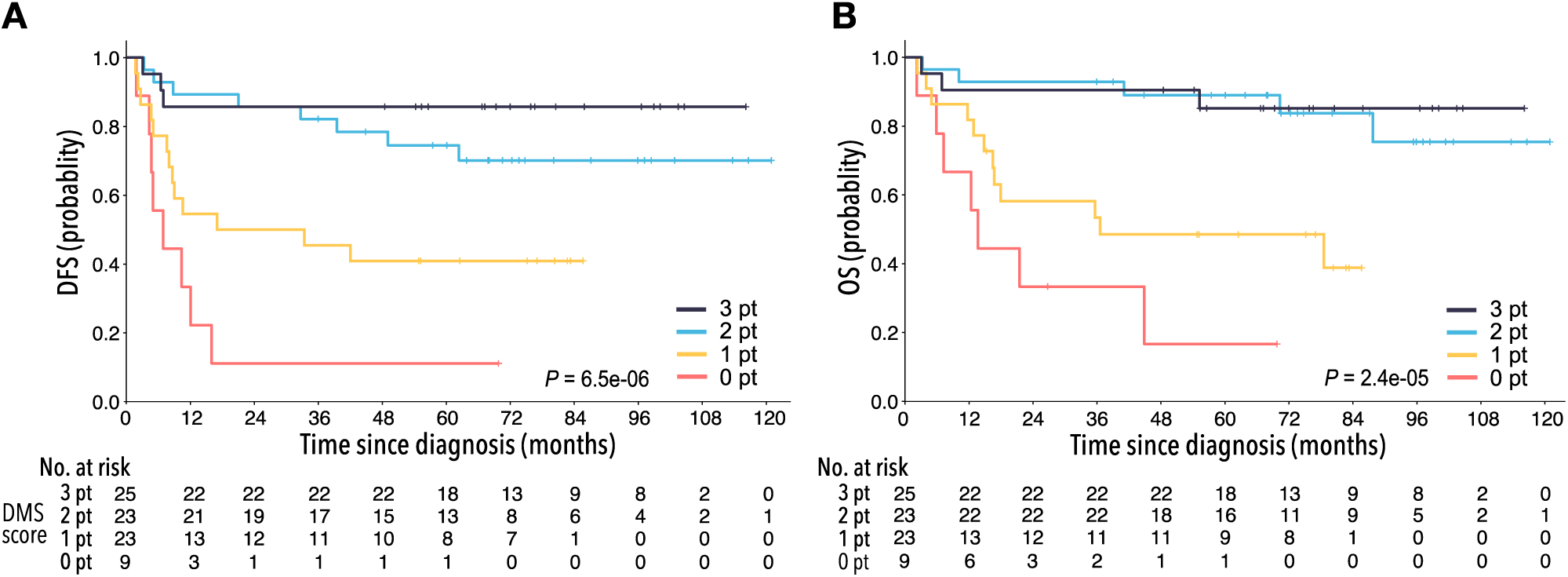
Validation of DMS score using independent cohorts. Kaplan-Meier analyses of (**A**) DFS and (**B**) OS based on DMS score using a validation cohort. A log-rank test was used for survival analysis.

### Multi-omics analysis reveals molecular background for prognostic impact of DMS

Finally, to test the impact of the GC-associated microenvironmental signature on genetic alterations serving as lymphoma cell hallmarks, we evaluated single nucleotide variants (SNVs), short insertion/deletions (indels) and copy number alterations (CNAs) in specimens from 106 patients with DLBCL. The most frequently mutated gene was KMT2D (MLL2), followed by PIM1, BCL6, BCL2, CDKN2A, and TP53 (fig. S7, A and B) in agreement with previous reports by us (30) and others (31–35). While ROS1 mutations were more frequent than previously reported, it is possible that some ROS1 SNVs are common variants unique to the Japanese population. In fact, eight of twelve ROS1 SNVs were defined as single nucleotide polymorphisms (SNPs) according to the Japanese Genome Variation Database (iJGVD). Fig. 6 summarizes the result of mutation analysis sorted by DMS score, plus GEP, COO classification, and “double-expressor” lymphoma (DEL) and “double-hit” lymphoma (DHL) status in 170 DLBCL patients of the training cohort. Expression of genes associated with poor prognosis, including cell cycle-related genes *FKBP4*, *MYC*, *PAICS*, *PDXP*, and *CDK4* (see also fig. S2), became higher as DMS score decreased (Fig. 6 upper).

**Fig. 6.**
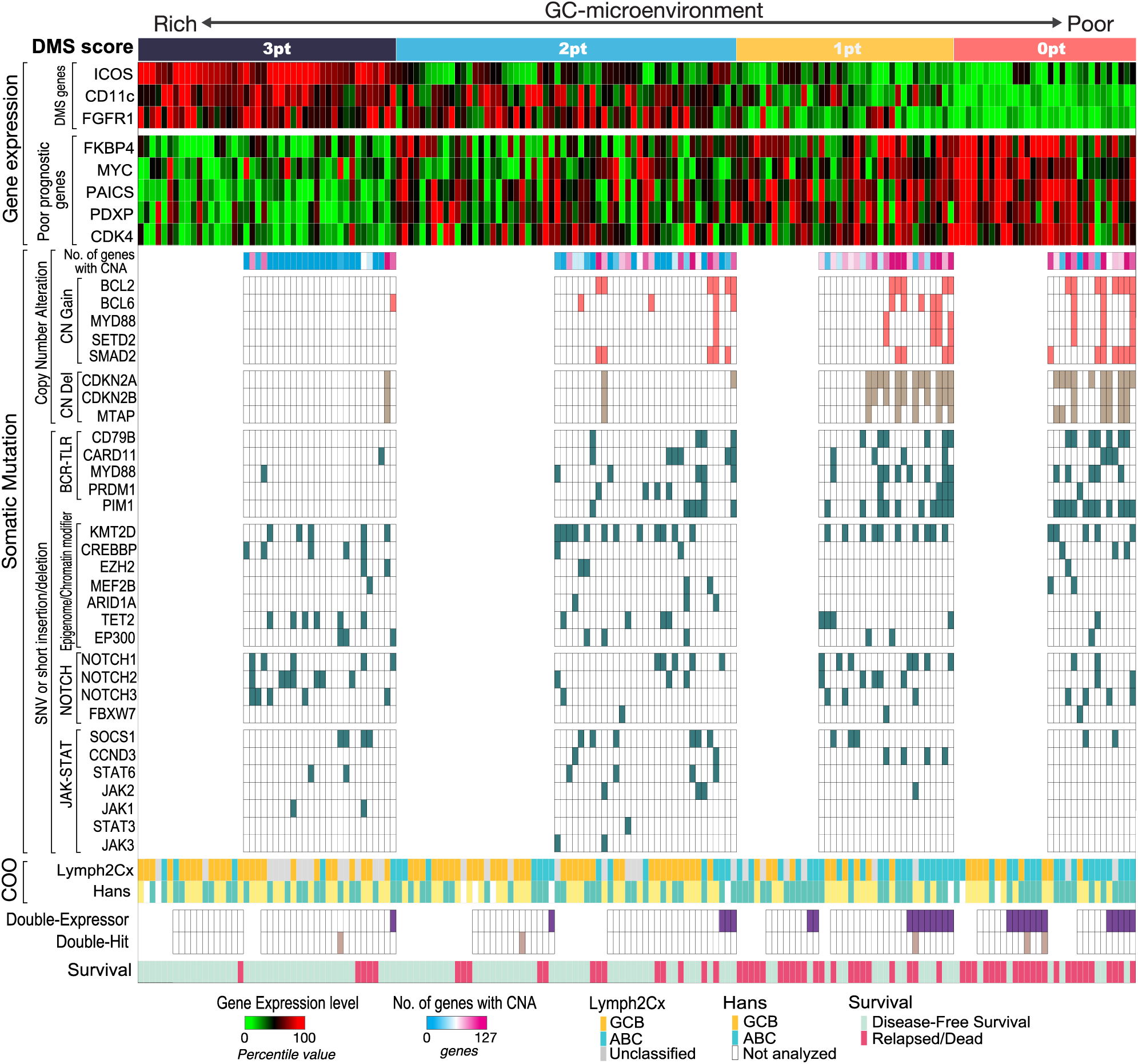
The relationship between the GC-microenvironment signature and multi-omics prognostic factors. ntegrated visualization of transcriptome data, genomic mutations, and COO classification combined with DMS score in 170 DLBCL cases. DMS scores were calculated based on enrichment of ICOS (follicular T cells), CD11c (DCs/macrophages), and FGFR1 (stromal cells). Tumor activity-related signatures, such as expression levels of proliferation-related genes or the number of genes showing CNA (copy number alteration) and BCR-TLR signaling mutations, were enriched as the DMS score decreased.

CNAs are dramatic structural changes in a genomic region (> 1kbp) and may involve multiple genes and alter global gene expression. In fact, CNA burden was more tightly associated with clinical outcomes than that of SNVs or short indels (fig. S7C). Expression of *ICOS*, *CD11c* or *FGFR1* marked patient groups with significantly fewer CNAs (fig. S7D). Likewise, cases with a DMS score of 3 were almost devoid of CNAs, and the overall CNA burden increased in low DMS cases (Fig. 6 middle and fig. S7D). CNAs related to poor prognosis such as CN gains of BCL2 (36) and CN deletions in the tumor suppressor CDKN2A and the adjacent MTAP gene at the chromosome 9p21 locus (37, 38) were also more frequent in low relative to high DMS cases (Fig. 6 middle). In addition, SNVs and short indels in BCR and Toll-like receptor (TLR) signaling genes and in the NF-kB regulator PIM1 (39) were more frequent in DMS-low cases, with statistical significance (Fig. 6 middle). These mutations as well as CN gains of MYD88 and adjacent SETD2 promote constitutive activation of anti-apoptotic and proliferative NF-kB pathways and contribute to poor prognosis (10, 30, 39–41). On the other hand, other major somatic mutations, including those involved in epigenome/chromatin modifiers, NOTCH- and JAK-STAT signaling pathways, were equally distributed among all DMS score groups.

We further assessed prognostic impact of the DMS score in DEL and DHL, both of which reportedly exhibit poor prognosis (42, 43). Among 123 DLBCL cases analyzed, 27 (22.0%; fig. S7, E and F) and 5 (4.1%; Fig. 6 lower) were identified as DEL and DHL, respectively. DEL were significantly enriched in DMS-low cases, and MYC protein levels inversely correlated with DMS score (fig. S4E), while DHL cases were not necessarily enriched in DMS-high cases likely due to the small sample size (Fig. 6 lower).

Thus, cell-intrinsic aggressiveness of DLBCL, which is characterized by proliferation-related gene expression, high frequency of genomic alterations, COO and DEL status, is associated with loss of GC-microenvironment components, as reflected by low DMS score. A GC-associated microenvironment comprehensively reflects multi-omics unfavorable molecular signatures.

## Discussion

GC formation is an essential step for normal B-cell development in LNs and requires recruitment of microenvironmental cells. Here, by employing accurate GEP, we demonstrated that GC-associated microenvironmental components have great impact on the clinical outcomes of DLBCL and established a DMS prognostic model. The DMS score successfully provided graded stratification of patients’ clinical responses to standard R-CHOP-based therapy. Moreover, DMS scoring had significant prognostic value even in patients’ groups stratified by either Lymph2Cx or the IPI score. These results suggest that interaction of lymphoma cells with their microenvironment governs DLBCL malignancy.

The nCounter system enabled highly sensitive, quantifiable analysis of fragmented RNAs in FFPE samples. In the original report proposing the COO stratification based on cDNA microarray analysis (1), microenvironmental cell-related transcripts were not extracted as prognostic factors. This omission may be due to the fact that conventional microarrays are not sensitive enough to quantify low abundance microenvironmental transcripts (44, 45). A previous study showed that microenvironmental cells function in DLBCL pathogenesis (46): gene expression profiling of DLBCL tissues revealed positive associations between extracellular matrix (ECM) deposition by stromal cells and favorable prognosis (“stromal-1” signature) and tumor blood-vessel density and poor prognosis (“stromal-2” signature) (46). The nCounter probe set used in our pilot study (Fig. 1A) included some of these genes, which were, in fact, prognostic, although no statistical significance was observed due presumably to small sample size (n=30) and/or differences in experimental systems (not shown). Accordingly, we identified stromal cell-related genes such as *FGFR1* and *COL8A2* as favorable prognostic indicators in this study (Fig. 1C).

Our nCounter data suggest follicular T cells and DC/macrophages as critical microenvironment components in determining DLBCL prognosis. In the normal reactive GC, follicular T cells patrol the B-cell region to participate in B-cell selection via the BCR, and their activity determines whether B-cells undergo apoptosis or proliferation (47, 48). Recent studies indicate two subsets of follicular T cells: Tfh cells expressing IL21, a cytokine promoting survival, proliferation and class switching of developing B-cells (49), and FOXP3-positive Tfr cells, which function in suppression and elimination of inappropriate B-cell clones (22). Tfr cells can negatively regulate Tfh cells by suppressing production of cytokines such as IL-21 or IL-4 and their proliferation (22). Our single cell analysis showed that both Tfh and Tfr cell populations infiltrated primary DLBCL tissues, although their ratios varied among specimens (Fig. 3). Tfh and Tfr cell function in DLBCL remains unclear, but the presence of follicular T cells is a strong indicator of favorable prognosis. In addition to follicular T cells, CD11c^+^ DC/macrophages play a critical role in clearing apoptotic B-cells (27) after selection by follicular T cells and participate in antigen-presentation to developing B-cells in normal GC. In our study, DC/macrophages were also enriched in primary DLBCL tissues from patients with favorable prognosis.

Since the DMS score was calculated based on the presence of three independent GC-associated microenvironmental cells, namely, stromal cells, follicular T cells, and DC/macrophages, these cells may cooperate to regulate DLBCL development or maintain DLBCL cells, comparable to their activity in the normal GC. Importantly, the DMS score stratified patients not only in cases of nodal DLBCL but also in those showing extranodal lesions (Fig. 4C), suggesting that in the latter the GC-associated microenvironment is reconstituted through recruitment of these cells into lymphoma tissue.

The nCounter probe set used in the pilot study broadly covers cancer and immune-related genes (1889 genes), enabling us to capture gene expression signatures associated with lymphoma microenvironment with high precision. Given its high sensitivity and reproducibility in measuring fragmented RNAs from FFPE samples, a more unbiased analysis targeting all transcripts, including non-coding RNAs, may further elucidate DLBCL biology relevant to clinical practice. Furthermore, while we believe that the simplicity of the DMS model is beneficial in terms of clinical utility, adding more genes, such as those positively-correlated with poor prognosis (such as *FKBP4* and *MYC*), may enhance the accuracy and reproducibility of the model.

Why does a high DMS score reflect favorable prognosis in DLBCL? One assumption is that lymphoma cells in favorable cases are regulated by GC-associated microenvironmental cells comparably to normally-developing B-cells, and once they acquire cell-autonomous proliferation and/or survival properties associated with malignancy, they become microenvironment-independent. This hypothesis is supported by the fact that DMS-low cases tend to accumulate lymphoma cell-intrinsic abnormalities, including expression of c-Myc-related or cell cycle-related molecules, and genomic alterations, all related to unfavorable prognosis (Fig. 6). In summary, our study shows that the GC-associated microenvironment signature is tightly associated with clinical outcomes in DLBCL patients. These data strongly suggest that evaluation of microenvironment components and their functions is the key to understand DLBCL pathogenesis and malignancy. The development of sophisticated quantitation methods focusing on microenvironment cells in combination with the COO and the IPI should be useful to predict DLBCL outcomes more accurately and to establish the best suitable treatment strategies.

## Materials and Methods

### Clinical data collection

We retrospectively collected de novo DLBCL cases, which were newly diagnosed by two or more experienced hematopathologists in the Kurume University Pathology Department from 2006 to 2013, based on the 2008 WHO classification of lymphoid neoplasms. Use of materials and clinical information was approved by the Research Ethics Committee of Kurume University in accordance with the Helsinki Declaration. FFPE samples, which were collected with clinical data, were anonymized before shipping to Kyushu University. The present study was approved by the institutional ethics committee of the Kyushu University Graduate School of Medical Sciences.

### GEP using nCounter system

Total RNA was extracted from DLBCL FFPE samples using a RNeasy FFPE extraction kit (QIAGEN, Hilden, Germany) after deparaffinization with Deparaffinization Solution (QIAGEN), based on the manufacturer’s instruction. RNA concentration was determined using a Qubit 2.0 Fluorometer (Thermo Fisher Scientific, Waltham, MA, USA). For analysis, 300 ng total RNA was hybridized to gene-specific probes at 65°C for 18 hours and then purified and deposited onto glass cartridges using the “high sensitivity” setting on the nCounter™ Prep Station (NanoString Technologies, Seattle, WA, USA). Gene expression levels were determined as the number of barcodes hybridized to specific RNA targets on cartridges, which were directly counted by the nCounter™ Digital Analyzer with 555 fields of view setting. Normalization of RNA loading was performed using the geometric mean of 15 housekeeping genes (ABCF1, ALAS1, EEF1G, G6PD, GAPDH, GUSB, HPRT1, OAZ1, POLR1B, POLR2A, PPIA, SDHA, TBP, TUBB, and RPL19). We utilized an nCounter Customer Assay Evaluation (CAE) kit to evaluate data reproducibility (Supplemental Figure 1A). To extract candidate prognostic genes in the screening cohort, we used PanCancer Pathways, Immunology, and Kinase panels. To validate the results of screening, we used a custom gene set (NanoString Technologies). In addition to DLBCL specimens, we profiled gene expression in the lymphoma lines: RC-K8, SU-DHL-1, SU-DHL-4, SU-DHL-9, Karpas-422 and OCI-Ly-3, as well as in five normal LN tissues. The latter were obtained by LN dissection as diagnosis for early stage gastric cancer and shown to be tumor cell-free. Detailed information about sample preparation and assay procedure is provided in Supplementary Methods.

### GEP by RNA sequencing

We removed rRNA from extracted total RNA using a Ribo-Zero rRNA Removal Kit (Epicentre, Madison, WI, USA) and then performed library preparation using a NEBNext Ultra Directional RNA Library Prep Kit for Illumina (New England BioLabs, Ipswich, MA, USA), according to the manufacturer’s instructions. Libraries generated were evaluated using KAPA Library Quantification Kits (KAPA Biosystems) and underwent 150 bp paired-end sequencing using the HiSeq 1500 system (Illumina, San Diego, CA, USA). Raw sequence reads were aligned to human reference hg19 using Hisat2 (50), and differential analysis was performed using HTSeq (51) and DESeq2 (52) to assess significance of gene expression changes.

### Single-cell GEP

CD4^+^ T cells were isolated from DLBCL tissues that had been cryopreserved in Cellbanker (TaKaRa Bio, Osaka, Japan). Cell sorting was performed using a BD FACS Aria III cell-sorting system (BD Biosciences, San Jose, CA, USA). Cells were sorted directly into Hank’s Balanced Salt solution (catalog number 14175-095, Thermo Fisher Scientific, Waltham, MA, USA) supplemented with 10% fetal bovine serum (Hyclone, Logan, UT, USA) and 2% EDTA (pH 8.0, Invitrogen, Carlsbad, CA, USA) and subjected to reverse transcription and pre-amplification with gene-specific primers using the C1 system (Fluidigm, South San Francisco, CA, USA), according to manufacturer’s instruction. Amplified cDNA was loaded onto a Fluidigm Dynamic Array Integrated Fluidic Chip and subjected to quantitative Real-Time PCR using a Biomark system (Fluidigm). Primers and probes were obtained from Applied Biosystems (Foster City, CA, USA). Cells exhibiting no B2M signals were omitted from the analysis.

### Multiplexed fluorescent immunostaining

Slides from biopsied tumor tissues were stained using an Opal multiplex tissue staining system (PerkinElmer, Waltham, MA, USA). Antigen retrieval was performed by heating slides to 93 ± 2 °C for 20 min in high-pH antigen unmasking solution (H-3301, Vector Labs, Burlingame, CA, USA), followed by blocking in 5% bovine serum albumin (Jackson ImmunoResearch, Birmingham, AL, USA) in PBS. Cell phenotyping and counting were performed using the Mantra quantitative pathology workstation (PerkinElmer) within representative fields pre-selected by trained hematopathologists. Spatial distribution and marker intensity in target cells were analyzed using inForm® image analysis (PerkinElmer) and Spotfire (TIBCO Software, Palo Alto, CA, USA) software.

### Calculation of the DMS score

The DMS score was calculated based on expression levels of three genes (*ICOS*, *CD11c* and *FGFR1*). To call positive (point=1) or negative (point=0) for each gene, the cutoff values were defined using the maximally selected rank statistics from “maxstat” R package. The DMS score was determined as a sum of points from each gene.

### Prognostic analysis

Data relevant to observation period and survival and/or disease status at the last observation was available for all study patients. OS was defined as the time from diagnosis to the last follow-up or death, and DFS as the time from diagnosis to any recurrence or death. Survival probability was estimated using the Kaplan–Meier method, and p-values determined by log-rank test. Univariate or multivariate analysis using a Cox proportional hazards model was used to assess the predictive value of DMS scores, as defined by gene expression analysis. Statistical analyses were performed using JMP Pro 13 software (SAS Institute Inc., Cary, NC, USA).

### Determination of cell-of-origin

DLBCL cell-of-origin subtypes were determined using Lymph2Cx and Hans criteria, according to previous reports (2, 28). Information relevant to cell-of-origin subtype based on Hans criteria was available for 160 of 170 DLBCL cases enrolled in the validation cohort.

### Mutation analysis

Among 170 cases in the training cohort, we randomly selected cases for mutational analysis and evaluated 120 FFPE-derived DNA samples. Genomic DNA was extracted using a QIAamp DNA FFPE Tissue Kit (QIAGEN) based on the manufacturer’s instruction. Then, we undertook massively parallel sequencing using a solution-phase SureSelect hybrid capture kit (Agilent Technologies, Santa Clara, CA, USA) and an HiSeq 2500 sequencer (Illumina). To detect genomic aberrations/rearrangements in FFPE samples, we used an OncoPanel assay (Brigham and Women’s Hospital) to evaluate exonic regions of 447 cancer genes and 191 regions across 60 genes (53). Among 120 cases, we analyzed data from 106 (14 samples were omitted due to poor data quality).

### Determination of DEL and DHL

Tissue microarrays (TMAs) were constructed from cores of 127 DLBCL tissues, whose FFPE samples were available from the training cohort. Among them, 122 cases were eligible for FISH and IHC analysis. Immunohistochemical staining was performed using antibodies against MYC (clone L-26; DakoCytomation, Glostrup, Denmark) and BCL2 (clone 124; DakoCytomation). Cutoff values for DEL were defined as described (42). FISH was performed using break-apart probes for MYC, BCL2 and BCL6, as described (54). Cases with split signals in ≥5% of nuclei were considered positive for the rearrangement. At least 100 cells per case were evaluated by two independent investigators (K.M. and H.M.).

### Analysis of publicly-available RNA-seq data sets

Two publicly-available data sets (10, 29) were used for validation. RNA-seq data from Schmitz et al study (29) were obtained from National Cancer Institute Genomic Data Commons (https://gdc.cancer.gov/about-data/publications/DLBCL-2018), and normalized FPKM values were used for survival analysis. The dataset from Reddy et al was obtained from the European Genome-phenome Archive (dataset id: EGAD00001003600), and processed and analyzed as described (55). The cutoff values were defined using receiver operating curve (ROC) analysis (56).

### Statistics

To compare data between two groups, we used the Wilcoxon Rank Sum Test for nonparametric testing. We used Pearson’s chi-square test for categorical data. Determination of cut-off values for gene expression levels were determined as the optimal value for predicting the relapse or death events by using the maximally selected rank statistics from the ‘maxstat’ R package. We set the cut-off values for three representative microenvironment genes, ICOS, CD11c and FGFR1, as 543.65, 2428.88, and 886.37, respectively. Univariate or multivariate analysis using a Cox proportional hazards model was used to assess the predictive value of prognostic models. Statistical analyses were performed using R (http://CRAN.R-project.org/) and JMP Pro 13 software (SAS Institute Inc., Cary, NC, USA).

## Supplementary Materials

Fig. S1. Accurate mRNA measurement enables the evaluation of tumor microenvironment.

Fig. S2. Clinical impact of extracted poor prognostic genes.

Fig. S3. ICOS, CD11c, and FGFR1 are representative genes for germinal center microenvironment.

Fig. S4. Prognostic impact of the DMS score.

Fig. S5. The DMS score was validated using two publicly-available DLBCL cohorts.

Fig. S6. The DMS score stratifies DLBCL patients independently of disease subtypes previously defined.

Fig. S7. Relationship between prognostic models and genomic mutations.

Table S1. Summary of Patient and pathology characteristics.

Table S2. Genes in Customer Assay Evaluation kit.

Table S3. Differentially Expressed Genes from Immunology Panel.

Table S4. Differentially Expressed Genes from Kinase Panel.

Table S5. Differentially Expressed Genes from PanCancer Pathway Panel.

Table S6. Differentially Expressed Genes from RNA-seq.

Table S7. Design and annotation of custom probe set for nCounter analysis.

Table S8. Multivariate analysis in the validation cohort.

Table S9. Multivariate analysis in Schmitz et al. NEJM 2018.

Table S10. Multivariate analysis in Reddy et al. Cell 2017.

## Acknowledgments

We thank all laboratory members and especially Drs. Masao Seto (Kurume University) and Shinya Rai (Kindai University) for valuable discussions; Dr. Joji Shimono for collecting clinical information; Drs. Hiroshi Arima, Momoko Nishikori and Akifumi Takaori-Kondo for the help on RNA-seq data analysis; and Dr. Elise Lamar for proofreading of manuscript.

## Author contributions

K.M., T. Maeda and K.A. designed experiments and wrote the manuscript. K.M., T.S., K.K., K.S., H.M., Y.S., Y. Kikushige, Y. Kunisaki, and F.K. performed experiments and analyzed data. T. Maeda, T. Miyamoto, H.I., J.A., Y.M., and K.O. reviewed the data.

