## Supplementary Materials for "A Germinal Center-Associated Microenvironmental Signature Reflects Malignant Phenotype and Outcome of Diffuse Large B-cell Lymphoma"

**Fig. S1. Accurate mRNA measurement enables the evaluation of tumor microenvironment.** (**A**) Reproducibility of mRNA quantification by the nCounter system. Scatter plot shows results of two analyses of 48 genes from the Microarray Quality Control Study (MAQC) of DLBCL tissues. Note that even low abundance genes were detected with high-reproducibility.
(**B**) A volcano plot indicates favorable (left) and poor (right) prognostic factors in the screening cohort. Note that microenvironmental cell-related genes were associated with favorable prognosis. A Mann-Whitney U test (unpaired) with bootstrap was performed. Differentially expressed genes with q<0.05 were shown.
(**C**) Shown are biopsy sites of DLBCL tissues analyzed in the validation cohort. Note that >40% of cases were extranodal.

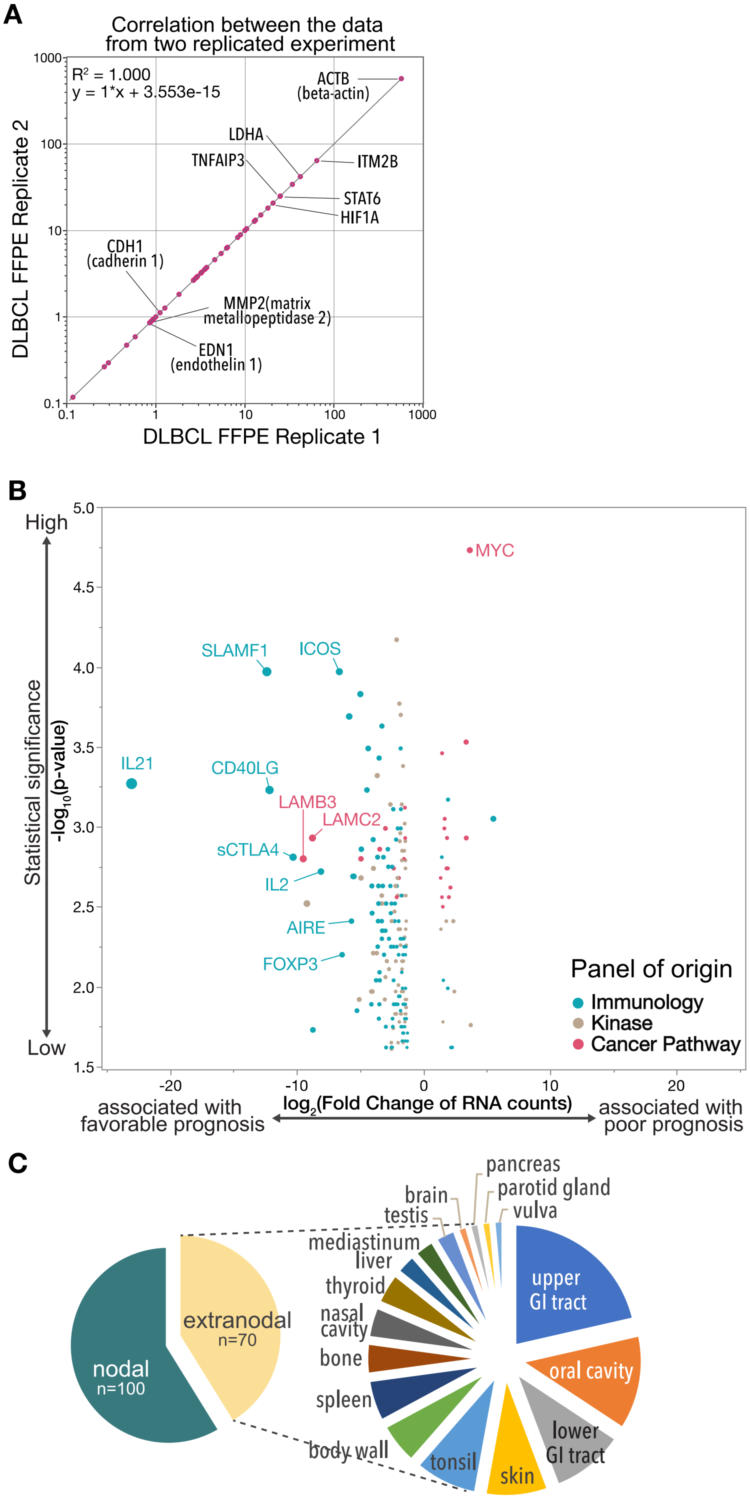

**Fig. S2. Clinical impact of extracted poor prognostic genes.** (**A**) Expression levels of four genes associated with unfavorable prognosis in lymphoma cell lines (n=6), primary DLBCL specimens (n=170), and normal lymph nodes (LNs) (n=5), shown as Box and Whisker plot (dots indicate outliers). Lymphoma lines analyzed were RC-K8, SU-DHL-1, SU-DHL-4, SU-DHL-9, Karpas-422, and OCI-Ly-3. All four genes are markers of cell proliferation. P-values were calculated using Steel-Dwass test (*p<0.05, **p<0.005, and ***p<0.0005). (**B**) Graphs show positive correlations between levels of MYC protein (x-axis) and transcript levels of proliferation-related genes, including MYC itself (y-axis). P-values were calculated using Pearson's product-moment correlation coefficient. (**C**) Kaplan-Meier curves show duration of disease-free survival (DFS) based on expression level genes shown in (A). P-values were calculated by log-rank test.

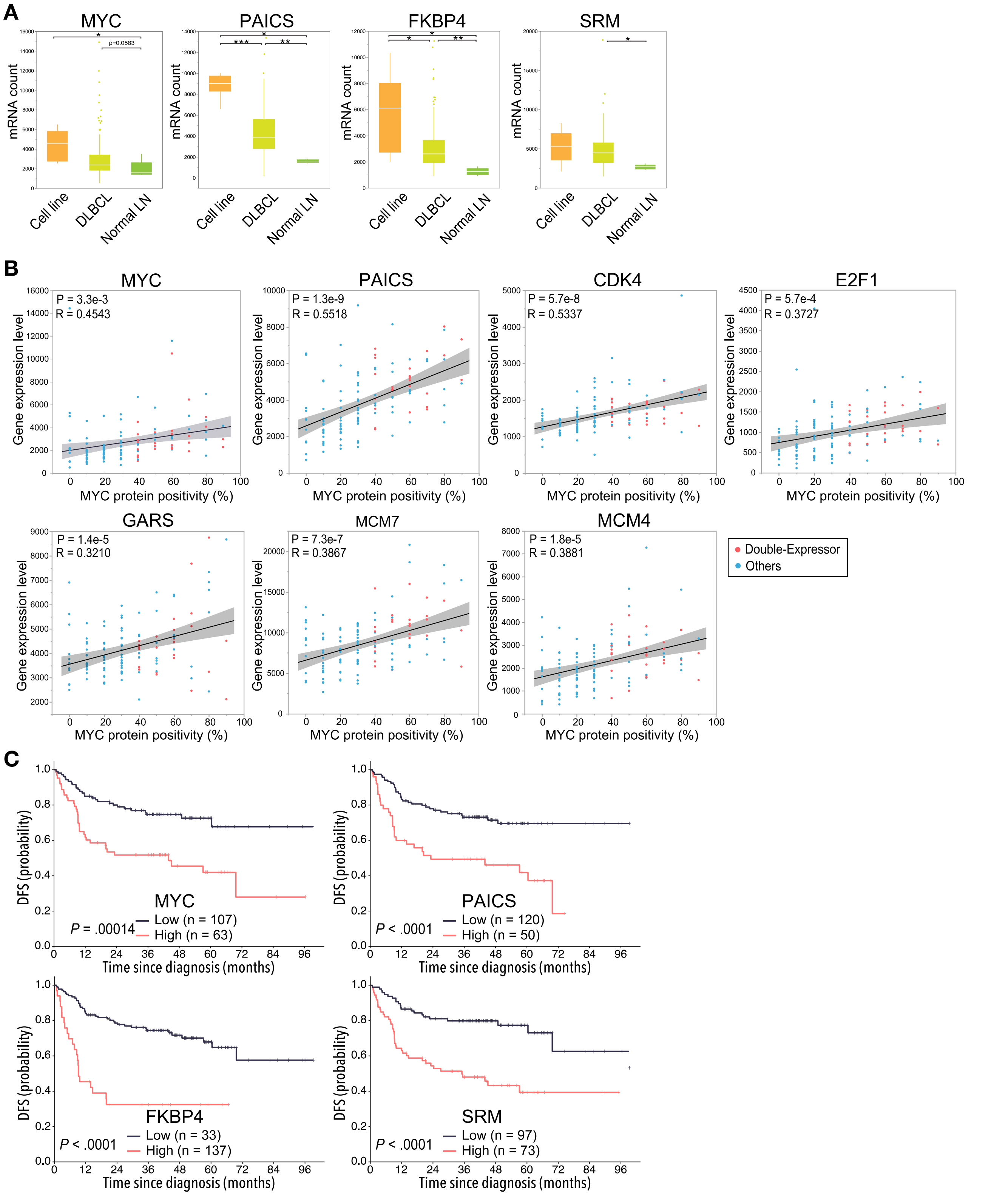

**Fig. S3. ICOS, CD11c, and FGFR1 are representative genes for germinal center microenvironment.** (**A**) Gene Ontology (GO) analysis of prognostic genes from the validation cohort. (**B**) Multispectral imaging shows steady-state lymph node (LN, upper) and reactive LN (lower) specimens stained with indicated markers. ICOS, CD11c and FGFR1 are enriched during germinal center (GC) formation. (**C**) Ratios of target ICOS+ follicular T cells (left) and CD11c+ DC/macrophages (Mø) (middle) to B-cells in follicle of steady-state LN and GC of reactive LN (delineated by dashed lines), as calculated by inForm image analysis software and compared by the Wilcoxon Rank Sum Test. Right panel shows ratio of the number of FGFR1-expressing cells to non-B/T/Mø cells in the follicle and GC.

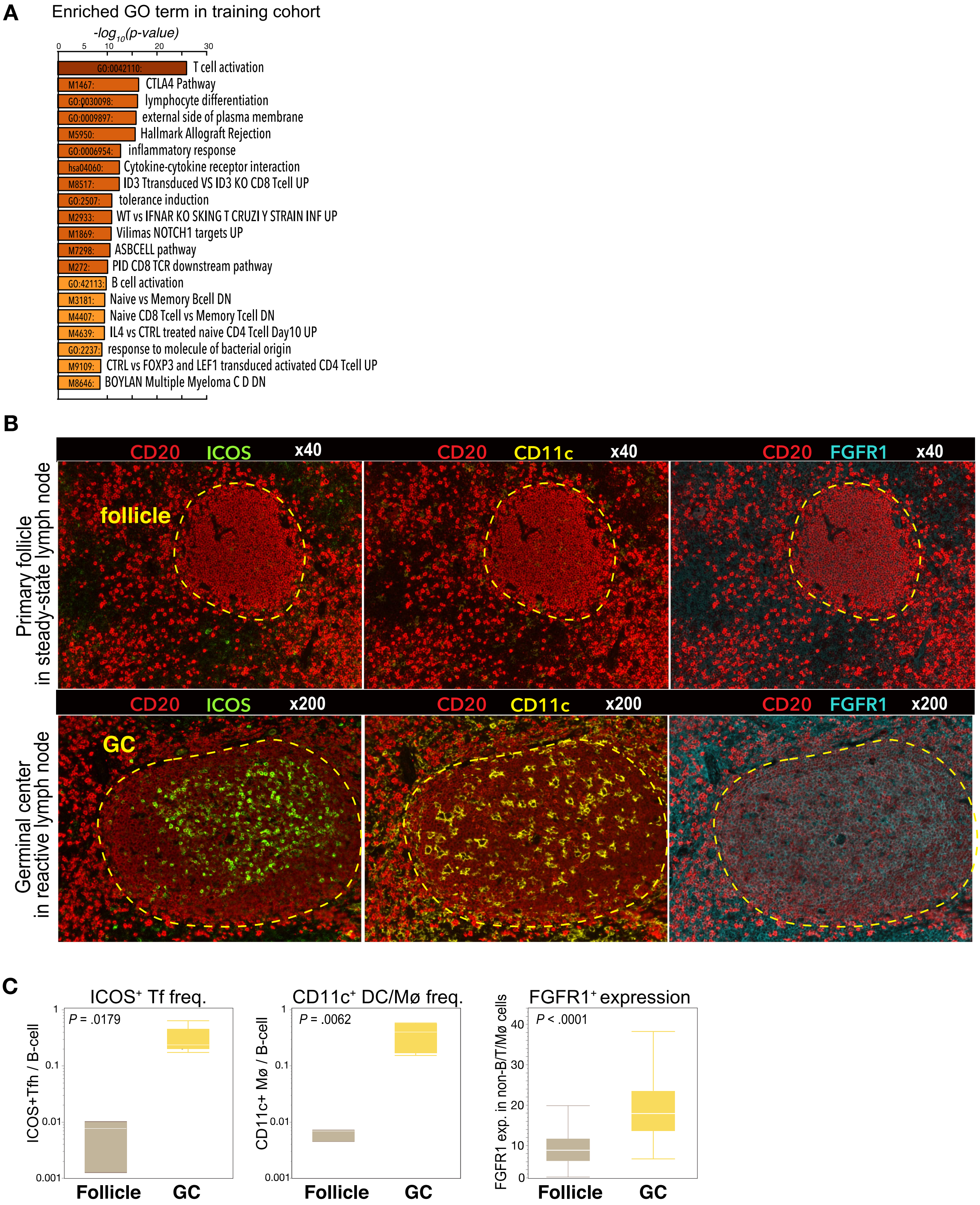

**Fig. S4. Prognostic impact of the DMS score.** (**A)** Kaplan-Meier overall survival (OS) curves based on the DMS score in all cases (left) and extranodal cases (right). P-values were calculated using the log-rank test. (**B**) mRNA levels of B cell-related genes in a given sample were comparable regardless of DMS score, suggesting well-balanced tumor content among samples. The Steel-Dwass multiple comparison test was performed.

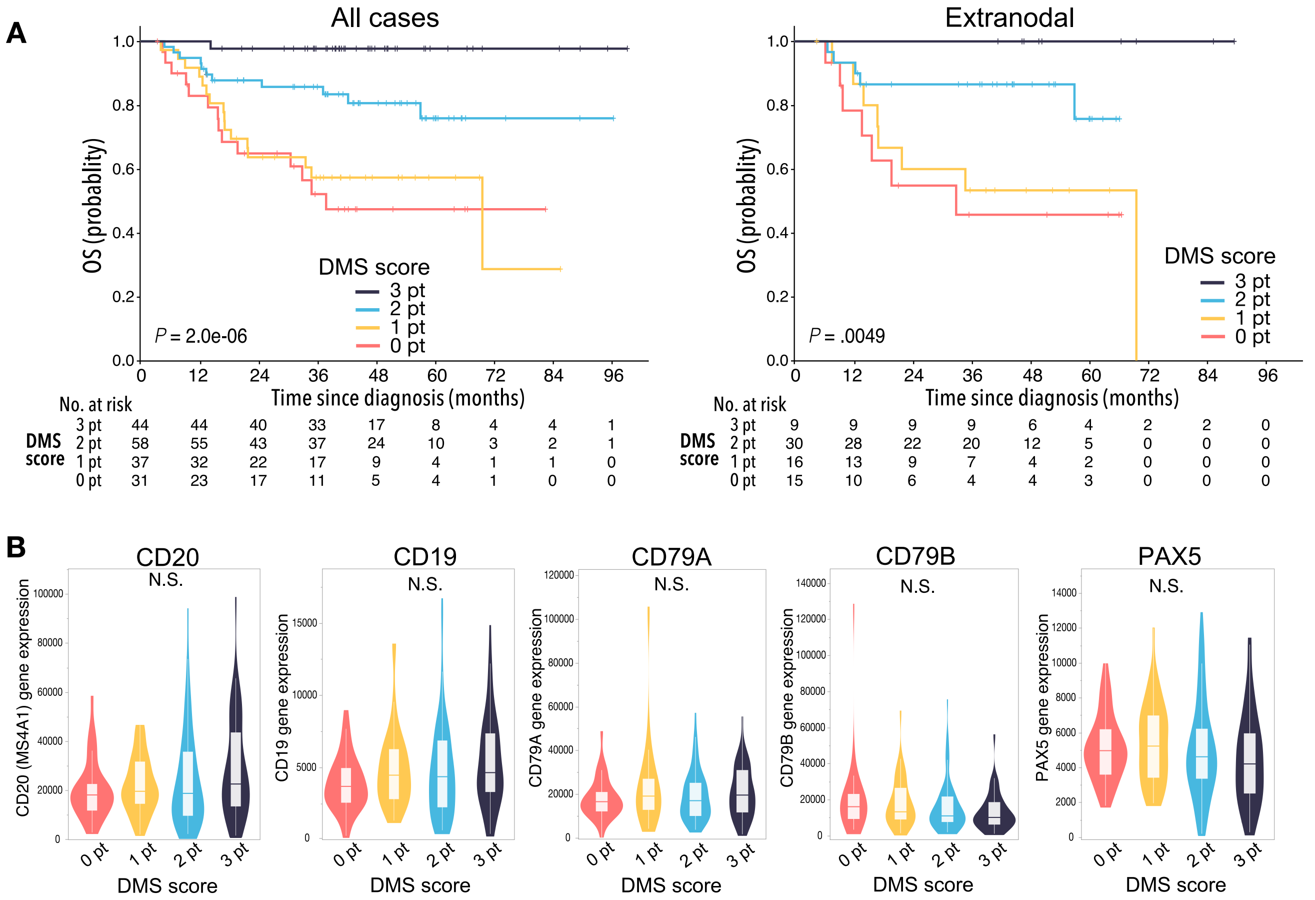


| DMS score (All cases) | 0 | 12 | 24 | 36 | 48 | 60 | 72 | 84 | 96 |
|---|---|---|---|---|---|---|---|---|---|
| 3 pt | 44 | 44 | 40 | 33 | 17 | 8 | 4 | 4 | 1 |
| 2 pt | 58 | 55 | 43 | 37 | 24 | 10 | 3 | 2 | 1 |
| 1 pt | 37 | 32 | 22 | 17 | 9 | 4 | 1 | 1 | 0 |
| 0 pt | 31 | 23 | 17 | 11 | 5 | 4 | 1 | 0 | 0 |

| DMS score (Extranodal) | 0 | 12 | 24 | 36 | 48 | 60 | 72 | 84 | 96 |
|---|---|---|---|---|---|---|---|---|---|
| 3 pt | 9 | 9 | 9 | 9 | 6 | 4 | 2 | 2 | 0 |
| 2 pt | 30 | 28 | 22 | 20 | 12 | 5 | 0 | 0 | 0 |
| 1 pt | 16 | 13 | 9 | 7 | 4 | 2 | 0 | 0 | 0 |
| 0 pt | 15 | 10 | 6 | 4 | 4 | 3 | 0 | 0 | 0 |

**Fig. S5. The DMS score was validated using two publicly-available DLBCL cohorts.** (**A**) Kaplan-Meier curves show duration of progression-free survival (PFS) in the Schmitz et al. cohort (top) and overall survival (OS) in the Reddy et al. cohort (bottom) based on expression levels of indicated genes. P-values were calculated by log-rank test. (**B**) Kaplan-Meier analyses of PFS (Schmitz cohort, left) and OS (Reddy cohort, right) based on DMS score. A log-rank test was used for survival analysis.

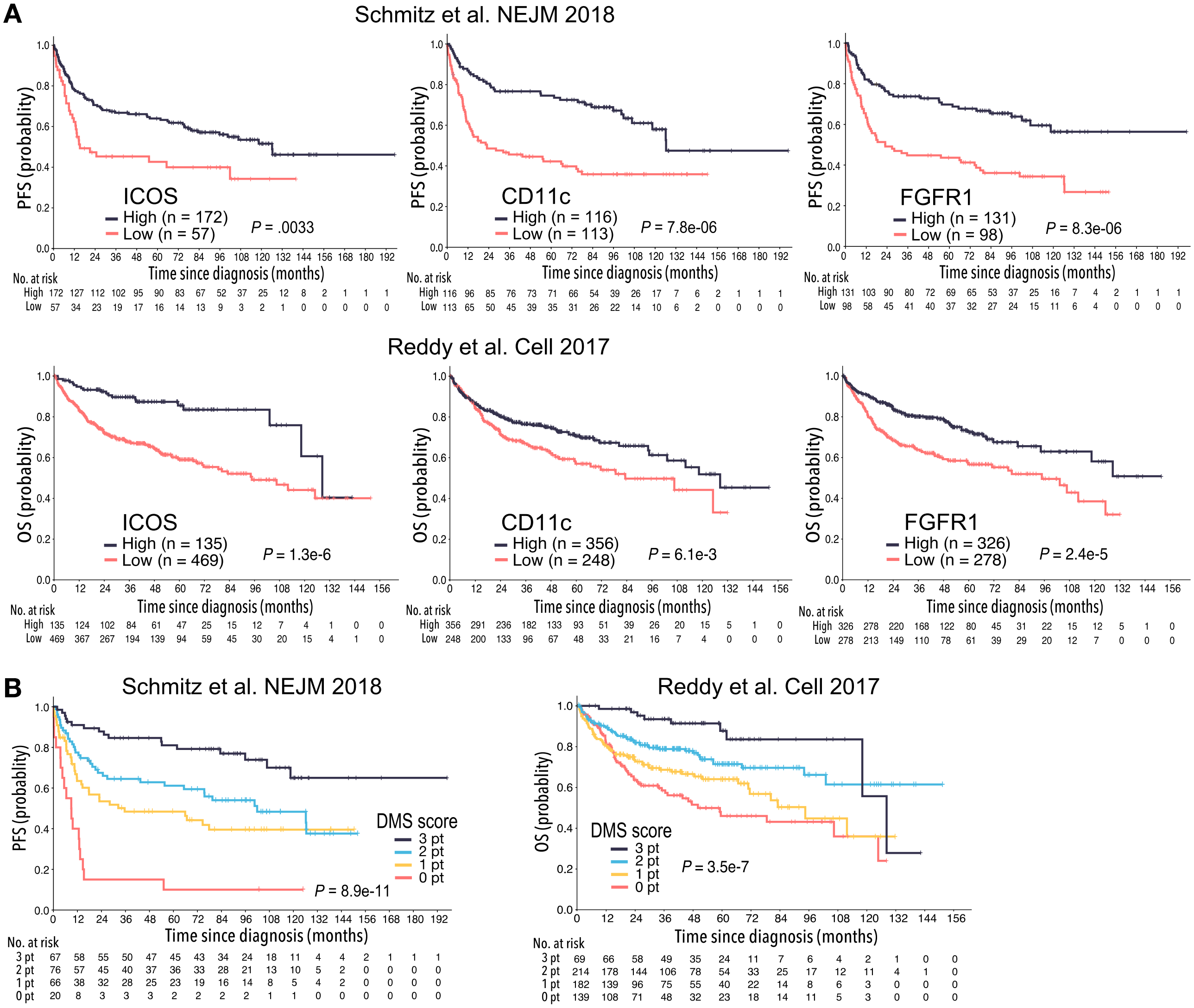

**Fig. S6. The DMS score stratifies DLBCL patients independently of disease subtypes previously defined.** Kaplan-Meier curves show duration of PFS or OS in disease subtypes reported by Schmitz et al (**A**) and Reddy et al (**B**). P-values were calculated by log-rank test.

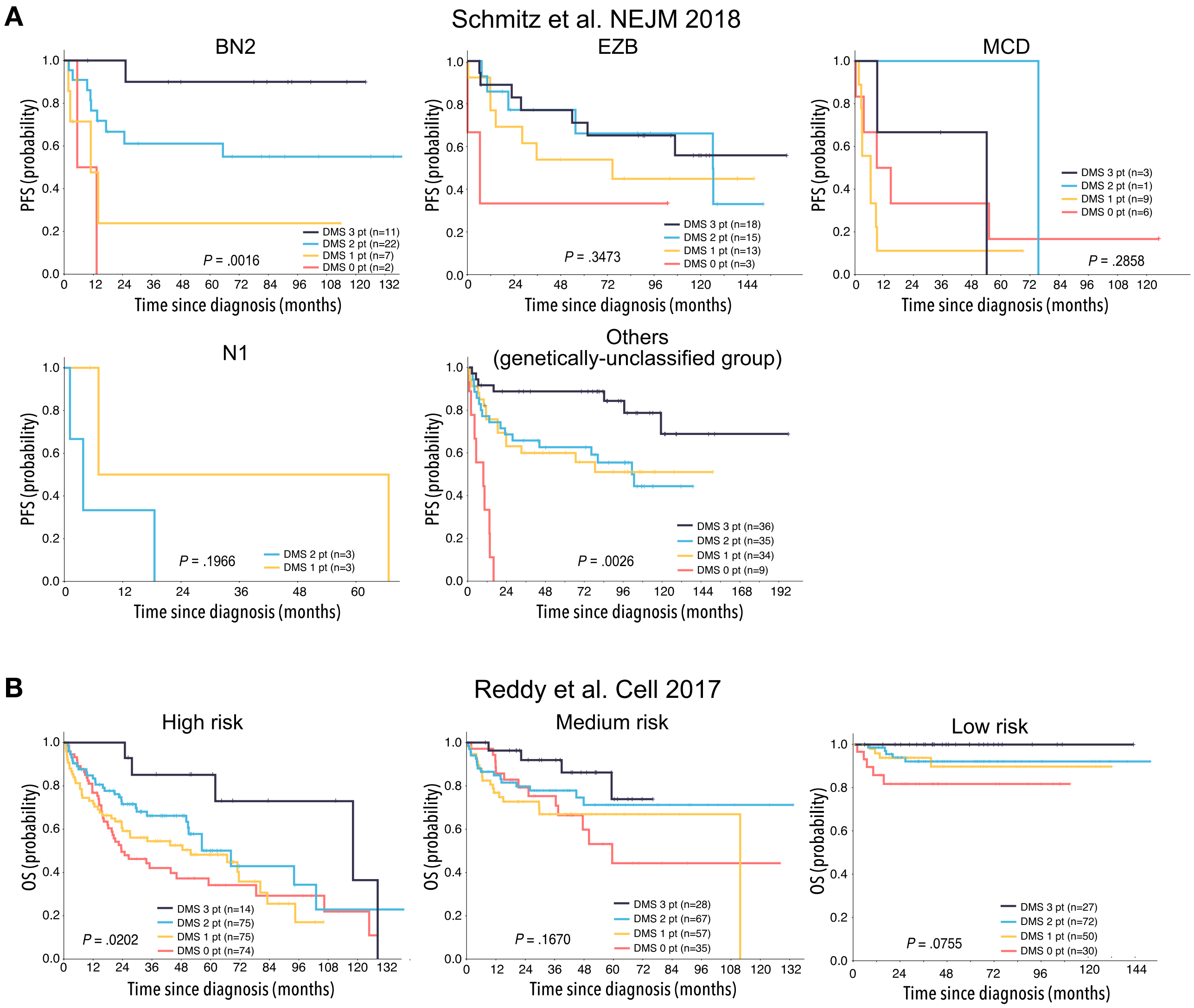


**Fig. S7. Relationship between prognostic models and genomic mutations.** (**A**, **B**) Bar graphs show frequency of cases with mutations (single nucleotide variants (SNVs), short insertion/deletions (indels), and copy number alterations (CNAs)) in specific genes in 106 patient specimens, as identified by the OncoPanel assay, and their distribution based on DMS score (**A**) and Lymph2Cx classification (**B**). (**C**) Prognostic value of mutation burden depending on indicated type of somatic mutation. Overall CNA burden had more significant prognostic impact than did the presence of SNVs or short indels. P-values were calculated by log-rank test. (**D**) Plots show differences in overall number of genes harboring CNAs based on DMS score or enrichment of indicated genes. ICOS$^{+}$ follicular T cell-rich cases were rarely affected by copy number changes. (**E**) MYC and BCL2 protein levels were assessed by immunohistochemistry. Dot graph shows MYC and BCL2 positivity in each sample (n=123). Cases positive for both MYC and BCL2 were defined as Double-expressor lymphoma (DEL). We call “positive” when more than 40% or 50% of the cells in a given section are positive for MYC or BCL2, respectively (depicted by a blue square). (**F**) DELs (magenta) exhibited low DMS scores. Fisher’s exact test was performed to calculate P-value. Numbers of cases are also shown.

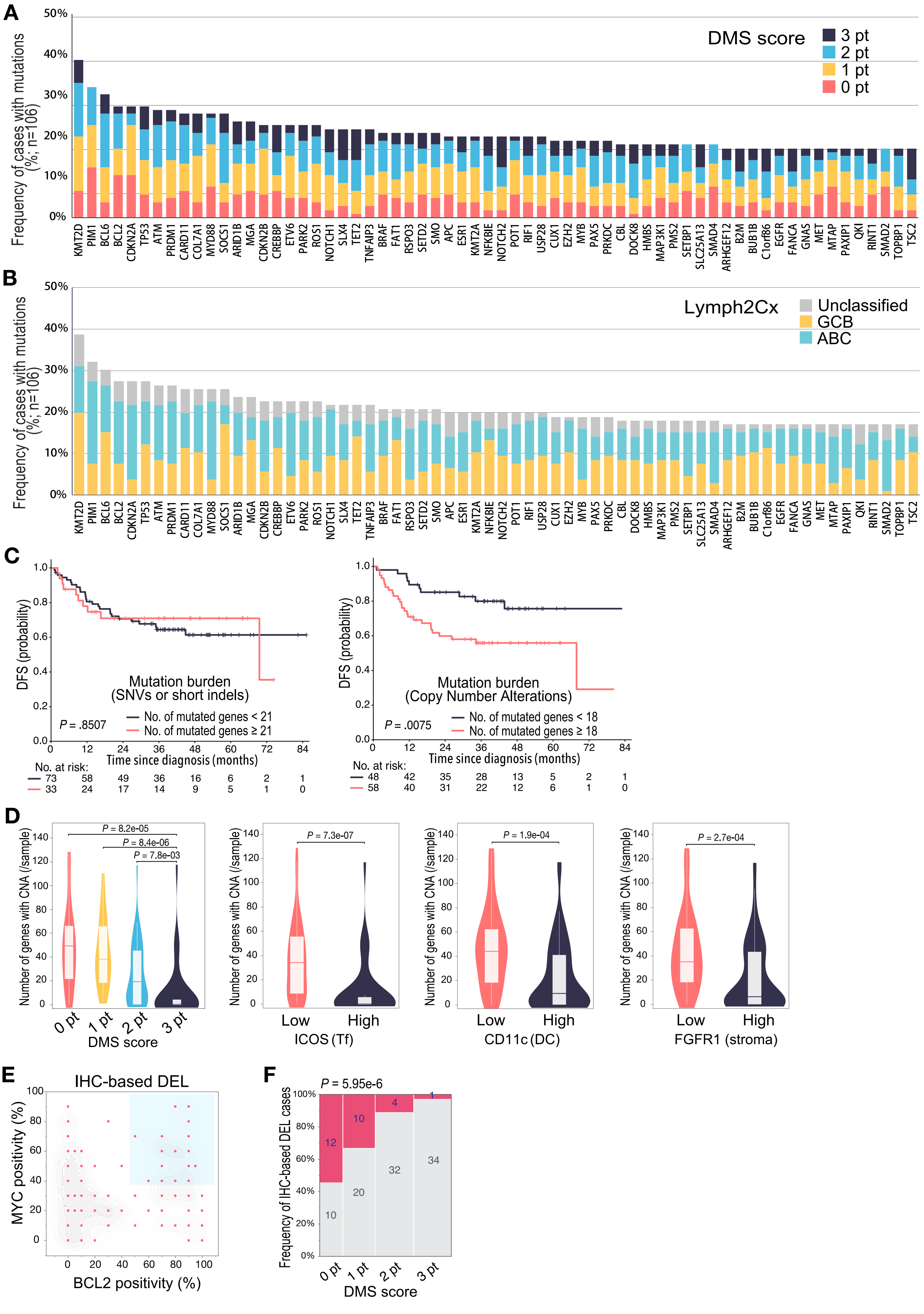

A
DMS score
3 pt
2 pt
1 pt
0 pt
Frequency of cases with mutations (%; n=106)
KMT2D PIM1 BCL6 BCL2 CDKN2A TP53 ATM PRDM1 CARD11 COL7A1 MYD88 SOCS1 ARID1B MGA CDKN2B CREBBP ETV6 PARK2 ROS1 NOTCH1 SLX4 TET2 TNFAIP3 BRAF FAT1 RSPO3 SETD2 SMO APC ESR1 KMT2A NFKBIE NOTCH2 POT1 RIF1 USP28 CUX1 EZH2 MYB PAX5 PRKDC CBL DOCK8 HMBS MAP3K1 PMS2 SETBP1 SLC25A13 SMAD4 ARHGEF12 B2M BUB1B C1orf86 EGFR FANCA GNAS MET MTAP PAXIP1 QKI RINT1 SMAD2 TOPBP1 TSC2
B
Lymph2Cx
Unclassified
GCB
ABC
C
DFS (probability)
Mutation burden (SNVs or short indels)
P = .8507
No. of mutated genes < 21
No. of mutated genes ≥ 21
Time since diagnosis (months)
No. at risk:
73 58 49 36 16 6 2 1
33 24 17 14 9 5 1 0
Mutation burden (Copy Number Alterations)
P = .0075
No. of mutated genes < 18
No. of mutated genes ≥ 18
48 42 35 28 13 5 2 1
58 40 31 22 12 6 1 0
D
Number of genes with CNA (/sample)
P = 8.2e-05
P = 8.4e-06
P = 7.8e-03
0 pt 1 pt 2 pt 3 pt
DMS score
P = 7.3e-07
Low High
ICOS (Tf)
P = 1.9e-04
CD11c (DC)
P = 2.7e-04
FGFR1 (stroma)
E
IHC-based DEL
MYC positivity (%)
BCL2 positivity (%)
F
P = 5.95e-6
Frequency of IHC-based DEL cases
12 10 10 20 4 32 1 34
DMS score

### Table S1. Summary of Patient and pathology characteristics

| Characteristic | Pilot cohort (n=30) | Training set (n=170) | Validation set (n=80) |
|---|---|---|---|
| Median age (range), years | 72.5 (36-83) | 71 (23-89) | 69 (18-88) |
| **Sex** | | | |
| Male | 20 (66.7) | 98 (57.6) | 49 (61.3) |
| Female | 10 (33.3) | 72 (42.4) | 31 (38.8) |
| **IPI risk group** | | | |
| Low | 10 (33.3%) | 49 (32.2%) | 32 (40.0%) |
| Low-intermediate | 4 (13.3%) | 29 (19.0%) | 13 (16.3%) |
| High-intermediate | 8 (26.7%) | 48 (31.6%) | 19 (23.8%) |
| High | 8 (26.7%) | 26 (17.1%) | 16 (20.0%) |
| **Initial treatment** | | | |
| R-CHOP | 24 (80.0%) | 119 (70.0%) | 61 (76.3%) |
| R-THPCOP | 6 (20.0%) | 51 (30.0%) | 19 (23.8%) |
| **Site of Biopsy** | | | |
| LN | 21 (70.0%%) | 100 (58.8%) | 50 (62.5%) |
| ExLN | 9 (30.0%%) | 70 (41.2%) | 30 (37.5%) |
| **Cell-of-origin subtype** | **Hans criteria** | **Lymph2Cx** | **Lymph2Cx** |
| GCB | 16 (53.3%) | 80 (47.0%) | 36 (45.0%) |
| ABC (non-GC) | 14 (46.7%) | 59 (34.7%) | 30 (37.5%) |
| Unclassified | | 31 (18.2%) | 14 (17.5%) |

### Table S2. Genes in Customer Assay Evaluation kit

| Official Symbol | Accession | Alias / Prev Symbol | Official Full Name |
|---|---|---|---|
| ABCC4 | NM_005845.2 | | ATP-binding cassette, sub-family C (CFTR/MRP), member 4 |
| ACTB | NM_001101.2 | | actin, beta |
| ADM | NM_001124.1 | | adrenomedullin |
| AMD1 | NM_001634.4 | | adenosylmethionine decarboxylase 1 |
| APC | NM_000038.3 | | adenomatous polyposis coli |
| ASPA | NM_000049.2 | | aspartoacylase (Canavan disease) |
| BTBD15 | NM_014155.2 | HSPC063, ZNF851 | zinc finger and BTB domain containing 44 |
| C11orf58 | NM_014267.3 | | chromosome 11 open reading frame 58 |
| C13orf23 | NM_025138.2 | bA50D16.2, FLJ12661 | chromosome 13 open reading frame 23 |
| CCNA2 | NM_001237.2 | | cyclin A2 |
| CDH1 | NM_004360.2 | UVO, uvomorulin, CD324 | cadherin 1, type 1, E-cadherin (epithelial) |
| CHGB | NM_001819.1 | | chromogranin B (secretogranin 1) |
| CYR61 | NM_001554.3 | IGFBP10, GIG1, CCN1 | cysteine-rich, angiogenic inducer, 61 |
| DNAJB9 | NM_012328.1 | | DnaJ (Hsp40) homolog, subfamily B, member 9 |
| EDN1 | NM_001955.2 | | endothelin 1 |
| EXOC2 | NM_018303.4 | SEC5L1, FLJ11026, Sec5p | exocyst complex component 2 |
| HBEGF | NM_001945.1 | | heparin-binding EGF-like growth factor |
| HDAC1 | NM_004964.2 | RPD3L1, HD1, GON-10 | histone deacetylase 1 |
| HIF1A | NM_001530.2 | | hypoxia inducible factor 1, alpha subunit (basic helix-loop-helix transcription factor) |
| HIST1H1D | NM_005320.2 | H1F3, H1.3 | histone cluster 1, H1d |
| ICAM1 | NM_000201.1 | | intercellular adhesion molecule 1 |
| IGF2R | NM_000876.1 | | insulin-like growth factor 2 receptor |
| IL1B | NM_000576.2 | | interleukin 1, beta |
| ITM2B | NM_021999.2 | | integral membrane protein 2B |
| KLF10 | NM_005655.1 | TIEG, EGRA, TIEG1 | Kruppel-like factor 10 |
| LDHA | NM_005566.1 | | lactate dehydrogenase A |
| LGI1 | NM_005097.1 | EPT, IB1099, ETL1, EPITEMPIN | leucine-rich, glioma inactivated 1 |
| MMP2 | NM_004530.2 | CLG4, CLG4A, TBE-1 | matrix metallopeptidase 2 (gelatinase A, 72kDa gelatinase, 72kDa type IV collagenase) |
| NDUFV3 | NM_021075.3 | | NADH dehydrogenase (ubiquinone) flavoprotein 3, 10kDa |
| NTS | NM_006183.3 | | neurotensin |
| POLR1B | NM_019014.3 | | polymerase (RNA) I polypeptide B, 128kDa |
| PPFIA2 | NM_003625.2 | | protein tyrosine phosphatase, receptor type, f polypeptide (PTPRF), interacting protein (liprin), alpha 2 |

| | | | |
|---|---|---|---|
| PSMC4 | NM_006503.2 | MIP224, TBP7, S6, MGC8570, MGC13687, MGC23214 | proteasome (prosome, macropain) 26S subunit, ATPase, 4 |
| PTGS2 | NM_000963.1 | | prostaglandin-endoperoxide synthase 2 (prostaglandin G/H synthase and cyclooxygenase) |
| RNF10 | NM_014868.3 | | ring finger protein 10 |
| SEC24D | NM_014822.1 | | SEC24 family, member D (S. cerevisiae) |
| SFRS10 | NM_004593.1 | Htra2-beta | transformer 2 beta homolog (Drosophila) |
| SHCBP1 | NM_024745.2 | | SHC SH2-domain binding protein 1 |
| STAT6 | NM_003153.3 | | signal transducer and activator of transcription 6, interleukin-4 induced |
| TDG | NM_003211.3 | | thymine-DNA glycosylase |
| TFRC | NM_003234.1 | | transferrin receptor (p90, CD71) |
| THBS1 | NM_003246.2 | | thrombospondin 1 |
| TNFAIP3 | NM_006290.2 | | tumor necrosis factor, alpha-induced protein 3 |
| TP53 | NM_000546.2 | p53, LFS1 | tumor protein p53 |
| TRAF4 | NM_004295.2 | | TNF receptor-associated factor 4 |
| TYMS | NM_001071.1 | TS, Tsase, TMS, HsT422 | thymidylate synthetase |
| UBE2B | NM_003337.2 | | ubiquitin-conjugating enzyme E2B (RAD6 homolog) |
| ZNF434 | NM_017810.2 | | zinc finger protein 434 |

### Table S3. Differentially Expressed Genes from Immunology Panel

| *Symbols* | *p-value* | *q-value* | *Regulation (Poor/Favorable)* | *Fold Change (Poor/Favorable)* |
|---|---|---|---|---|
| ICOS | 1.07E-04 | 1.04E-02 | down | -6.64 |
| SLAMF1 | 1.07E-04 | 1.04E-02 | down | -12.36 |
| CD96 | 1.48E-04 | 1.04E-02 | down | -4.98 |
| CR2 | 2.03E-04 | 1.04E-02 | down | -5.86 |
| EGR2 | 2.37E-04 | 1.04E-02 | down | -3.28 |
| CD6 | 3.22E-04 | 1.04E-02 | down | -4.36 |
| IKBKE | 3.22E-04 | 1.04E-02 | down | -1.80 |
| DUSP4 | 3.74E-04 | 1.05E-02 | down | -3.50 |
| IL21 | 5.39E-04 | 1.20E-02 | down | -23.04 |
| CD40LG | 5.84E-04 | 1.20E-02 | down | -12.15 |
| CD7 | 5.84E-04 | 1.20E-02 | down | -4.46 |
| CD3EAP | 6.75E-04 | 1.26E-02 | up | 1.94 |
| CD46 | 7.80E-04 | 1.26E-02 | down | -1.78 |
| LCP2 | 7.80E-04 | 1.26E-02 | down | -2.38 |
| ICAM5 | 8.97E-04 | 1.35E-02 | up | 5.52 |
| MALT1 | 1.03E-03 | 1.37E-02 | down | -2.03 |
| NFATC2 | 1.03E-03 | 1.37E-02 | down | -1.79 |
| IL12RB1 | 1.19E-03 | 1.41E-02 | down | -2.16 |
| ZAP70 | 1.19E-03 | 1.41E-02 | down | -3.97 |
| IKZF2 | 1.37E-03 | 1.41E-02 | down | -2.81 |
| TCF7 | 1.37E-03 | 1.41E-02 | down | -4.90 |
| sCTLA4 | 1.56E-03 | 1.41E-02 | down | -10.29 |
| CD28 | 1.57E-03 | 1.41E-02 | down | -3.16 |
| CD3D | 1.57E-03 | 1.41E-02 | down | -3.62 |
| POLR1B | 1.57E-03 | 1.41E-02 | up | 1.45 |
| FYN | 1.79E-03 | 1.46E-02 | down | -2.77 |
| NFKB2 | 1.79E-03 | 1.46E-02 | down | -2.47 |
| IL2 | 1.89E-03 | 1.46E-02 | down | -8.09 |
| CTLA4-TM | 2.05E-03 | 1.46E-02 | down | -5.49 |
| CTLA4_all | 2.05E-03 | 1.46E-02 | down | -5.56 |
| CD5 | 2.34E-03 | 1.46E-02 | down | -3.55 |
| IL21R | 2.34E-03 | 1.46E-02 | down | -2.10 |
| ITGAL | 2.34E-03 | 1.46E-02 | down | -2.13 |
| JAK2 | 2.34E-03 | 1.46E-02 | down | -2.61 |
| SH2D1A | 2.34E-03 | 1.46E-02 | down | -4.07 |
| STAT4 | 2.34E-03 | 1.46E-02 | down | -3.21 |
| PTPRC_all | 2.66E-03 | 1.62E-02 | down | -1.74 |

| | | | | |
|---|---|---|---|---|
| CD2 | 3.03E-03 | 1.67E-02 | down | -3.55 |
| CD3E | 3.03E-03 | 1.67E-02 | down | -2.95 |
| CD58 | 3.03E-03 | 1.67E-02 | down | -2.52 |
| IL2RG | 3.03E-03 | 1.67E-02 | down | -2.25 |
| CD45R0 | 3.44E-03 | 1.81E-02 | down | -2.58 |
| TNFRSF4 | 3.44E-03 | 1.81E-02 | down | -4.05 |
| AIRE | 3.87E-03 | 1.84E-02 | down | -5.68 |
| ITGAX | 3.91E-03 | 1.84E-02 | down | -3.58 |
| PTPN6 | 3.91E-03 | 1.84E-02 | down | -1.96 |
| TIGIT | 3.91E-03 | 1.84E-02 | down | -3.29 |
| TMEM173 | 3.91E-03 | 1.84E-02 | down | -2.28 |
| BATF3 | 4.43E-03 | 1.96E-02 | down | -3.21 |
| CD80 | 4.43E-03 | 1.96E-02 | down | -3.00 |
| TRAF1 | 4.43E-03 | 1.96E-02 | down | -3.26 |
| FAS | 5.01E-03 | 2.02E-02 | down | -2.72 |
| IL2RB | 5.01E-03 | 2.02E-02 | down | -3.28 |
| ITGA4 | 5.01E-03 | 2.02E-02 | down | -1.82 |
| ITGB2 | 5.01E-03 | 2.02E-02 | down | -1.68 |
| TRAF5 | 5.01E-03 | 2.02E-02 | down | -1.55 |
| CIITA | 5.66E-03 | 2.06E-02 | down | -2.11 |
| CXCR3 | 5.66E-03 | 2.06E-02 | down | -3.61 |
| IL6R | 5.66E-03 | 2.06E-02 | down | -2.56 |
| JAK3 | 5.66E-03 | 2.06E-02 | down | -2.36 |
| MCL1 | 5.66E-03 | 2.06E-02 | down | -1.53 |
| ZEB1 | 5.66E-03 | 2.06E-02 | down | -1.39 |
| BCL10 | 6.38E-03 | 2.15E-02 | down | -1.61 |
| CCBP2 | 6.38E-03 | 2.15E-02 | down | -3.11 |
| FOXP3 | 6.38E-03 | 2.15E-02 | down | -6.42 |
| IKBKG | 6.38E-03 | 2.15E-02 | down | -1.43 |
| SLAMF6 | 6.38E-03 | 2.15E-02 | down | -1.96 |
| CD86 | 8.09E-03 | 2.64E-02 | down | -1.71 |
| LIF | 8.09E-03 | 2.64E-02 | down | -3.47 |
| C1QBP | 9.08E-03 | 2.81E-02 | up | 1.57 |
| KLRB1 | 9.08E-03 | 2.81E-02 | down | -3.74 |
| MS4A1 | 9.08E-03 | 2.81E-02 | down | -2.36 |
| TNFRSF9 | 9.08E-03 | 2.81E-02 | down | -3.45 |
| CD48 | 1.02E-02 | 3.02E-02 | down | -1.65 |
| SLC2A1 | 1.02E-02 | 3.02E-02 | up | 1.92 |
| TGFBR1 | 1.02E-02 | 3.02E-02 | down | -1.41 |
| CD19 | 1.14E-02 | 3.26E-02 | down | -1.95 |
| ICAM3 | 1.14E-02 | 3.26E-02 | down | -1.91 |

| | | | | |
|---|---|---|---|---|
| NOTCH1 | 1.14E-02 | 3.26E-02 | down | -1.86 |
| CD8B | 1.28E-02 | 3.27E-02 | down | -3.48 |
| IL11RA | 1.28E-02 | 3.27E-02 | down | -2.35 |
| MAF | 1.28E-02 | 3.27E-02 | down | -2.25 |
| NFATC3 | 1.28E-02 | 3.27E-02 | down | -1.42 |
| PAX5 | 1.28E-02 | 3.27E-02 | down | -1.61 |
| PDCD1 | 1.28E-02 | 3.27E-02 | down | -4.11 |
| RELB | 1.28E-02 | 3.27E-02 | down | -1.51 |
| SOCS1 | 1.28E-02 | 3.27E-02 | down | -2.03 |
| VCAM1 | 1.28E-02 | 3.27E-02 | down | -2.11 |
| CXCL13 | 1.43E-02 | 3.57E-02 | down | -5.25 |
| ICOSLG | 1.43E-02 | 3.57E-02 | down | -1.81 |
| MAP4K4 | 1.59E-02 | 3.81E-02 | down | -1.48 |
| NOD2 | 1.59E-02 | 3.81E-02 | down | -2.44 |
| STAT5B | 1.59E-02 | 3.81E-02 | down | -1.34 |
| TBX21 | 1.59E-02 | 3.81E-02 | down | -3.52 |
| C3 | 1.77E-02 | 3.99E-02 | down | -2.97 |
| IFI35 | 1.77E-02 | 3.99E-02 | down | -1.77 |
| IFNGR1 | 1.77E-02 | 3.99E-02 | down | -1.66 |
| IL7R | 1.77E-02 | 3.99E-02 | down | -2.42 |
| PLAU | 1.77E-02 | 3.99E-02 | down | -1.72 |
| TRAF3 | 1.77E-02 | 3.99E-02 | down | -1.48 |
| CCL22 | 1.87E-02 | 4.16E-02 | down | -8.73 |
| IKBKB | 1.97E-02 | 4.27E-02 | down | -1.63 |
| JAK1 | 1.97E-02 | 4.27E-02 | down | -1.56 |
| RELA | 1.97E-02 | 4.27E-02 | down | -1.26 |
| ATG16L1 | 2.19E-02 | 4.57E-02 | down | -1.26 |
| IFNAR2 | 2.19E-02 | 4.57E-02 | down | -1.61 |
| NFKBIA | 2.19E-02 | 4.57E-02 | down | -1.68 |
| STAT2 | 2.19E-02 | 4.57E-02 | down | -1.43 |
| ATG12 | 2.43E-02 | 4.80E-02 | down | -1.30 |
| BTLA | 2.43E-02 | 4.80E-02 | down | -2.95 |
| CD163 | 2.43E-02 | 4.80E-02 | up | 2.17 |
| GPR183 | 2.43E-02 | 4.80E-02 | down | -2.47 |
| HLA-DRB1 | 2.43E-02 | 4.80E-02 | down | -2.53 |
| S100A9 | 2.43E-02 | 4.80E-02 | up | 2.27 |

### Table S4. Differentially Expressed Genes from Kinase Panel

| *Symbols* | *p-value* | *q-value* | *Regulation (Poor/Favorable)* | *Fold Change (Poor/Favorable)* |
|---|---|---|---|---|
| MAST3 | 6.78E-05 | 1.22E-02 | down | -2.12 |
| STK17B | 1.72E-04 | 1.22E-02 | down | -1.90 |
| PRKD2 | 2.00E-04 | 1.22E-02 | down | -1.79 |
| IKBKE | 4.14E-04 | 1.75E-02 | down | -1.61 |
| ITK | 4.78E-04 | 1.75E-02 | down | -3.65 |
| FYN | 7.25E-04 | 1.89E-02 | down | -2.61 |
| RPS6KA3 | 7.25E-04 | 1.89E-02 | down | -1.71 |
| CSNK1D | 9.52E-04 | 1.89E-02 | down | -1.48 |
| STK17A | 1.09E-03 | 1.89E-02 | down | -1.83 |
| BMP2K | 1.24E-03 | 1.89E-02 | down | -1.87 |
| PDK3 | 1.24E-03 | 1.89E-02 | down | -1.43 |
| CDK17 | 1.42E-03 | 1.89E-02 | down | -1.52 |
| LIMK1 | 1.42E-03 | 1.89E-02 | down | -1.68 |
| CDK13 | 1.61E-03 | 1.89E-02 | down | -1.42 |
| PTK2B | 1.61E-03 | 1.89E-02 | down | -1.70 |
| ALPK2 | 1.84E-03 | 1.89E-02 | down | -3.95 |
| SCYL2 | 1.84E-03 | 1.89E-02 | down | -1.42 |
| HUNK | 2.08E-03 | 1.89E-02 | down | -4.93 |
| FER | 2.09E-03 | 1.89E-02 | down | -1.95 |
| SGK1 | 2.09E-03 | 1.89E-02 | down | -2.05 |
| FGFR1 | 2.37E-03 | 1.89E-02 | down | -2.21 |
| NEK6 | 2.37E-03 | 1.89E-02 | down | -2.18 |
| EPHA3 | 2.68E-03 | 1.89E-02 | down | -3.29 |
| GRK5 | 2.68E-03 | 1.89E-02 | down | -2.04 |
| SCYL3 | 2.68E-03 | 1.89E-02 | down | -1.66 |
| TAF1 | 2.68E-03 | 1.89E-02 | down | -1.39 |
| CAMK2B | 3.03E-03 | 1.99E-02 | down | -9.20 |
| EPHA1 | 3.03E-03 | 1.99E-02 | down | -3.28 |
| OBSCN | 3.43E-03 | 2.15E-02 | down | -2.28 |
| CDKL2 | 3.87E-03 | 2.15E-02 | down | -2.51 |
| NRBP1 | 3.87E-03 | 2.15E-02 | down | -1.36 |
| GSG2 | 3.87E-03 | 2.15E-02 | up | 1.80 |
| MAPK12 | 3.87E-03 | 2.15E-02 | up | 2.36 |
| DDR2 | 4.36E-03 | 2.16E-02 | down | -1.96 |
| NPR2 | 4.36E-03 | 2.16E-02 | down | -1.84 |
| STK24 | 4.36E-03 | 2.16E-02 | down | -1.40 |
| CDK4 | 4.36E-03 | 2.16E-02 | up | 1.38 |
| CDK19 | 4.90E-03 | 2.37E-02 | down | -1.48 |
| MAP2K4 | 5.51E-03 | 2.53E-02 | down | -1.36 |
| PRKACB | 5.51E-03 | 2.53E-02 | down | -1.81 |

| | | | | |
|---|---|---|---|---|
| BRSK1 | 6.19E-03 | 2.71E-02 | down | -3.96 |
| STK32B | 6.19E-03 | 2.71E-02 | down | -3.69 |
| DAPK2 | 6.93E-03 | 2.71E-02 | down | -1.70 |
| DSTYK | 6.93E-03 | 2.71E-02 | down | -1.58 |
| IKBKB | 6.93E-03 | 2.71E-02 | down | -1.64 |
| JAK3 | 6.93E-03 | 2.71E-02 | down | -2.18 |
| ROR2 | 6.93E-03 | 2.71E-02 | down | -2.89 |
| CLK1 | 7.76E-03 | 2.74E-02 | down | -1.69 |
| MAP3K8 | 7.76E-03 | 2.74E-02 | down | -1.69 |
| TNIK | 7.76E-03 | 2.74E-02 | down | -2.72 |
| ZAP70 | 8.68E-03 | 3.01E-02 | down | -3.00 |
| PRKD1 | 9.69E-03 | 3.25E-02 | down | -1.97 |
| ACVR1 | 1.08E-02 | 3.25E-02 | down | -1.59 |
| EPHB6 | 1.08E-02 | 3.25E-02 | down | -4.12 |
| JAK1 | 1.08E-02 | 3.25E-02 | down | -1.58 |
| MARK1 | 1.08E-02 | 3.25E-02 | down | -4.03 |
| TLK1 | 1.08E-02 | 3.25E-02 | down | -1.73 |
| TRIO | 1.08E-02 | 3.25E-02 | down | -2.19 |
| DYRK3 | 1.08E-02 | 3.25E-02 | up | 2.44 |
| AURKA | 1.20E-02 | 3.30E-02 | down | -1.41 |
| BRAF | 1.20E-02 | 3.30E-02 | down | -1.33 |
| CAMK2G | 1.20E-02 | 3.30E-02 | down | -1.43 |
| MUSK | 1.20E-02 | 3.30E-02 | down | -5.07 |
| TXK | 1.20E-02 | 3.30E-02 | down | -3.05 |
| AKT1 | 1.34E-02 | 3.46E-02 | down | -1.35 |
| ILK | 1.34E-02 | 3.46E-02 | down | -1.40 |
| MAPK3 | 1.34E-02 | 3.46E-02 | down | -1.34 |
| ALPK1 | 1.49E-02 | 3.46E-02 | down | -1.80 |
| CDC42BPA | 1.49E-02 | 3.46E-02 | down | -1.75 |
| JAK2 | 1.49E-02 | 3.46E-02 | down | -2.14 |
| PRKCH | 1.49E-02 | 3.46E-02 | down | -2.24 |
| ROCK2 | 1.49E-02 | 3.46E-02 | down | -1.38 |
| TGFBR1 | 1.49E-02 | 3.46E-02 | down | -1.45 |
| AKT3 | 1.65E-02 | 3.48E-02 | down | -1.91 |
| MAP4K4 | 1.65E-02 | 3.48E-02 | down | -1.46 |
| PBK | 1.65E-02 | 3.48E-02 | up | 1.54 |
| CAMKV | 1.73E-02 | 3.62E-02 | up | 3.73 |
| CDK14 | 1.83E-02 | 3.77E-02 | down | -2.38 |
| CASK | 2.23E-02 | 4.23E-02 | down | -2.00 |
| MST1R | 2.23E-02 | 4.23E-02 | down | -2.54 |
| YES1 | 2.23E-02 | 4.23E-02 | down | -1.61 |
| DCLK1 | 2.46E-02 | 4.53E-02 | down | -2.52 |

### Table S5. Differentially Expressed Genes from PanCancer Pathway Panel

| *Symbols* | *p-value* | *q-value* | *Regulation (Poor/Favorable)* | *Fold Change (Poor/Favorable)* |
|---|---|---|---|---|
| MYC | 1.85.E-05 | 6.98.E-03 | up | 3.66 |
| MAPK12 | 2.98.E-04 | 3.81.E-02 | up | 3.37 |
| POLR2H | 3.50.E-04 | 3.81.E-02 | up | 1.47 |
| TTC31 | 7.59.E-04 | 3.81.E-02 | down | -1.46 |
| CDK4 | 8.82.E-04 | 3.81.E-02 | up | 1.68 |
| DUSP4 | 1.02.E-03 | 3.81.E-02 | down | -3.00 |
| MCM7 | 1.02.E-03 | 3.81.E-02 | up | 1.65 |
| LAMC2 | 1.18.E-03 | 3.81.E-02 | down | -8.76 |
| LAMC3 | 1.18.E-03 | 3.81.E-02 | up | 3.38 |
| MCM4 | 1.18.E-03 | 3.81.E-02 | up | 1.86 |
| ERCC6 | 1.18.E-03 | 3.81.E-02 | down | -1.44 |
| CC2D1B | 1.37.E-03 | 3.81.E-02 | down | -3.45 |
| LAMB3 | 1.58.E-03 | 3.81.E-02 | down | -9.49 |
| MYCN | 1.58.E-03 | 3.81.E-02 | down | -4.93 |
| CDC14A | 1.58.E-03 | 3.81.E-02 | down | -1.54 |
| FAS | 1.82.E-03 | 3.81.E-02 | down | -2.35 |
| ENDOG | 1.82.E-03 | 3.81.E-02 | up | 1.92 |
| MFNG | 1.82.E-03 | 3.81.E-02 | up | 1.82 |
| FGFR1 | 2.10.E-03 | 3.94.E-02 | down | -1.94 |
| MAD2L2 | 2.10.E-03 | 3.94.E-02 | up | 1.36 |
| CDKN2C | 2.41.E-03 | 4.32.E-02 | up | 2.13 |
| DLL4 | 2.76.E-03 | 4.33.E-02 | down | -2.11 |
| E2F1 | 2.76.E-03 | 4.33.E-02 | up | 2.01 |
| HDAC2 | 2.76.E-03 | 4.33.E-02 | up | 1.46 |
| CDC25B | 3.16.E-03 | 4.76.E-02 | up | 1.53 |

### Table S6. Differentially Expressed Genes from RNA-seq

| *Symbols* | *p-value* | *Regulation (Poor/Favorable)* | *Fold Change (Poor/Favorable)* |
|---|---|---|---|
| ICOS | 3.71.E-07 | down | -2.14 |
| SULF2 | 1.85.E-06 | up | 1.59 |
| TERT | 2.19.E-06 | up | 2.08 |
| ATRNL1 | 2.53.E-06 | down | -1.81 |
| B4GALT2 | 4.60.E-06 | up | 1.11 |
| SUMO4 | 5.77.E-06 | down | -1.07 |
| CCR4 | 9.55.E-06 | down | -1.87 |
| SNUPN | 1.19.E-05 | up | 0.63 |
| IFRD2 | 1.42.E-05 | up | 0.88 |
| PSAT1 | 1.52.E-05 | up | 1.50 |
| CD80 | 1.56.E-05 | down | -1.37 |
| MRTO4 | 2.32.E-05 | up | 0.95 |
| LYAR | 2.47.E-05 | up | 1.07 |
| PDE4D | 2.73.E-05 | down | -1.21 |
| TAB2 | 2.76.E-05 | down | -0.78 |
| CTLA4 | 2.89.E-05 | down | -1.72 |
| EGR2 | 3.07.E-05 | down | -1.37 |
| CD58 | 4.04.E-05 | down | -1.02 |
| TBC1D4 | 4.98.E-05 | down | -1.39 |
| MRPL46 | 5.17.E-05 | up | 0.90 |
| LRRC2 | 5.41.E-05 | down | -1.66 |
| FKBP4 | 6.61.E-05 | up | 1.21 |
| EIF3B | 7.21.E-05 | up | 0.70 |
| GRAMD1C | 7.24.E-05 | down | -1.55 |
| TRAF1 | 7.54.E-05 | down | -1.12 |
| FOXRED2 | 7.64.E-05 | up | 1.05 |
| C19orf10 | 7.74.E-05 | up | 0.87 |
| ATAD3A | 7.88.E-05 | up | 1.06 |
| NUDT4 | 8.11.E-05 | down | -1.34 |
| E2F2 | 1.08.E-04 | up | 1.04 |
| NOP14 | 1.22.E-04 | up | 0.65 |
| EPB41L4B | 1.28.E-04 | down | -1.77 |
| DPY19L2P2 | 1.36.E-04 | up | 1.00 |
| C1QBP | 1.42.E-04 | up | 1.00 |
| TRIM65 | 1.43.E-04 | up | 0.69 |
| SHMT2 | 1.46.E-04 | up | 0.97 |
| SARDH | 1.46.E-04 | down | -1.48 |
| IL21 | 1.53.E-04 | down | -1.83 |
| CD40LG | 1.59.E-04 | down | -1.81 |
| ACAT1 | 1.61.E-04 | up | 0.81 |

| | | | |
|---|---|---|---|
| FAS | 1.81.E-04 | down | -1.05 |
| ICA1 | 2.13.E-04 | down | -1.58 |
| TIMM8B | 2.14.E-04 | up | 0.68 |
| RAB27A | 2.47.E-04 | down | -0.63 |
| CD36 | 2.48.E-04 | up | 1.53 |
| MYC | 3.10.E-04 | up | 1.19 |
| BYSL | 3.17.E-04 | up | 0.75 |
| PDXP | 3.35.E-04 | up | 0.94 |
| MYBBP1A | 3.82.E-04 | up | 0.91 |
| CD96 | 3.85.E-04 | down | -0.97 |
| SRM | 4.37.E-04 | up | 0.89 |
| HIST1H3I | 4.43.E-04 | up | 1.22 |
| TFAP4 | 4.73.E-04 | up | 0.86 |
| CYP2U1 | 5.93.E-04 | up | 1.07 |
| XYLB | 6.17.E-04 | up | 0.90 |
| MEIS2 | 6.79.E-04 | up | 1.17 |
| GAR1 | 8.10.E-04 | up | 0.71 |
| PAICS | 9.19.E-04 | up | 0.93 |
| BZW2 | 9.20.E-04 | up | 0.72 |
| RABL2A | 1.02.E-03 | down | -0.86 |
| GARS | 1.05.E-03 | up | 0.57 |
| PSPH | 1.33.E-03 | up | 0.82 |
| ETFA | 1.35.E-03 | up | 0.53 |

### Table S7. Design and annotation of custom probe set for nCounter analysis

| Gene Identifier | Sources-1 | Sources-2 | Sources-3 | Accession | Position |
|---|---|---|---|---|---|
| ABCF1 | Additional_House_Keeping_Genes | | | NM_001090.2 | 851-950 |
| ACAT1 | RNAseq | | | NM_000019.3 | 1347-1446 |
| CCBP2 | nCounter_Immunology | | | NM_001296.3 | 1346-1445 |
| ACVR1 | nCounter_Kinase | | | NM_001105.2 | 1666-1765 |
| AGK | Additional_House_Keeping_Genes | | | NM_018238.3 | 817-916 |
| AIRE | nCounter_Immunology | | | NM_000383.2 | 1865-1964 |
| AKT1 | nCounter_Kinase | | | NM_001014432.1 | 1276-1375 |
| AKT3 | nCounter_Kinase | | | NM_005465.4 | 288-387 |
| ALAS1 | Additional_House_Keeping_Genes | | | NM_000688.4 | 396-495 |
| ALPK1 | nCounter_Kinase | | | NM_025144.2 | 2976-3075 |
| ALPK2 | nCounter_Kinase | | | NM_052947.3 | 6126-6225 |
| AMMECR1L | Additional_House_Keeping_Genes | | | NM_031445.2 | 253-352 |
| ARFGEF3 | Additional_Others | | | NM_020340.2 | 246-345 |
| ARG1 | Additional_Macrophage | | | NM_000045.3 | 11-110 |
| ASB13 | Additional_Lymph2Cx | | | NM_024701.3 | 1636-1735 |
| ATAD3A | RNAseq | | | NM_018188.2 | 626-725 |
| ATG12 | nCounter_Immunology | | | NM_004707.2 | 26-125 |
| ATG16L1 | nCounter_Immunology | | | NM_017974.3 | 2406-2505 |
| ATRNL1 | RNAseq | | | NR_074088.1 | 1459-1558 |
| AURKA | nCounter_Kinase | | | NM_003600.2 | 406-505 |
| B3GAT1 | Additional_T_cell_subset | | | NM_054025.2 | 1034-1133 |
| B4GALT2 | RNAseq | | | NM_001005417.2 | 1125-1224 |
| BATF3 | nCounter_Immunology | | | NM_018664.2 | 870-969 |
| BCL10 | nCounter_Immunology | | | NM_003921.2 | 1251-1350 |
| BCL2 | Additional_Recurrently_mutated_in_DLBCL | | | NM_000657.2 | 6-105 |
| BCL6 | Additional_T_cell_subset | | | NM_138931.1 | 506-605 |
| BMP2K | nCounter_Kinase | | | NM_017593.3 | 1256-1355 |
| BRAF | nCounter_Kinase | | | NM_004333.3 | 566-665 |
| BRSK1 | nCounter_Kinase | | | NM_032430.1 | 1056-1155 |
| BTK | Additional_BCR_signal | | | NM_000061.1 | 571-670 |
| BTLA | nCounter_Immunology | | | NM_181780.2 | 306-405 |
| BYSL | RNAseq | | | NM_004053.3 | 1081-1180 |
| BZW2 | RNAseq | | | NR_027624.1 | 837-936 |
| C1QBP | nCounter_Immunology | RNAseq | | NM_001212.3 | 746-845 |
| C3 | nCounter_Immunology | | | NM_000064.2 | 4397-4496 |
| CAMK2B | nCounter_Kinase | | | NM_001220.3 | 366-465 |
| CAMK2G | nCounter_Kinase | | | NM_001222.2 | 234-333 |

| | | | | | |
|---|---|---|---|---|---|
| CAMKV | nCounter_Kinase | | | NM_024046.3 | 631-730 |
| CARD11 | Additional_BCR_signal | | | NM_032415.2 | 1076-1175 |
| CASK | nCounter_Kinase | | | NM_003688.1 | 2046-2145 |
| CC2D1B | nCounter_PanCancer_Pathway | | | NM_032449.2 | 1387-1486 |
| CCDC50 | Additional_Lymph2Cx | | | NM_174908.3 | 919-1018 |
| CCL22 | nCounter_Immunology | | | NM_002990.3 | 798-897 |
| CCR3 | Additional_T_cell_subset | | | NM_001837.2 | 981-1080 |
| CCR4 | RNAseq | | | NM_005508.4 | 36-135 |
| CD163 | nCounter_Immunology | | | NM_004244.4 | 1631-1730 |
| CD19 | nCounter_Immunology | | | XM_011545981.1 | 714-813 |
| CD2 | nCounter_Immunology | | | NM_001767.2 | 1401-1500 |
| CD200 | Additional_T_cell_subset | | | NM_005944.5 | 453-552 |
| CD226 | Additional_Macrophage | | | NM_001303618.1 | 845-944 |
| CD27 | Additional_T_cell_subset | | | NM_001242.3 | 1201-1300 |
| CD28 | nCounter_Immunology | | | NM_001243078.1 | 2066-2165 |
| CD36 | RNAseq | | | NM_000072.3 | 708-807 |
| CD3D | nCounter_Immunology | | | NM_000732.4 | 111-210 |
| CD3E | nCounter_Immunology | | | NM_000733.2 | 76-175 |
| CD3EAP | nCounter_Immunology | | | NM_012099.1 | 556-655 |
| CD3G | Additional_T_cell_subset | | | NM_000073.2 | 405-504 |
| CD4 | Additional_T_cell_subset | | | NM_000616.4 | 976-1075 |
| CD40LG | nCounter_Immunology | RNAseq | | NM_000074.2 | 1226-1325 |
| CD46 | nCounter_Immunology | | | NM_172350.1 | 366-465 |
| CD48 | nCounter_Immunology | | | NM_001778.2 | 271-370 |
| CD5 | nCounter_Immunology | | | NM_014207.2 | 1296-1395 |
| CD58 | nCounter_Immunology | RNAseq | | NM_001779.2 | 479-578 |
| CD6 | nCounter_Immunology | | | NM_001254751.1 | 1723-1822 |
| CD7 | nCounter_Immunology | | | NM_006137.6 | 441-540 |
| CD79A | Additional_BCR_signal | | | NM_001783.3 | 696-795 |
| CD79B | Additional_BCR_signal | | | NM_021602.2 | 25-124 |
| CD80 | nCounter_Immunology | RNAseq | | NM_005191.3 | 675-774 |
| CD81 | Additional_Macrophage | | | NM_004356.3 | 1375-1474 |
| CD86 | nCounter_Immunology | | | NM_175862.3 | 1266-1365 |
| CD8B | nCounter_Immunology | | | NM_172099.2 | 440-539 |
| CD96 | nCounter_Immunology | RNAseq | | NM_005816.4 | 1135-1234 |
| CDC14A | nCounter_PanCancer_Pathway | | | NM_033313.2 | 1401-1500 |
| CDC25B | nCounter_PanCancer_Pathway | | | XR_937181.1 | 1543-1642 |
| CDC42BPA | nCounter_Kinase | | | NM_003607.2 | 7221-7320 |
| CDK13 | nCounter_Kinase | | | NM_003718.4 | 1535-1634 |
| CDK14 | nCounter_Kinase | | | NM_001287135.1 | 507-606 |
| CDK17 | nCounter_Kinase | | | NM_002595.2 | 1476-1575 |

| | | | | | |
|---|---|---|---|---|---|
| CDK19 | nCounter_Kinase | | | NM_015076.3 | 4106-4205 |
| CDK4 | nCounter_Kinase | nCounter_PanCancer_Pathway | | NM_000075.2 | 1056-1155 |
| CDKL2 | nCounter_Kinase | | | NM_003948.2 | 871-970 |
| CDKN2C | nCounter_PanCancer_Pathway | | | NM_001262.2 | 1296-1395 |
| CHUK | Additional_BCR_signal | | | NM_001278.3 | 861-960 |
| CIITA | nCounter_Immunology | | | NM_000246.3 | 471-570 |
| MGL1 | Additional_Macrophage | | | NM_182906.2 | 503-602 |
| CLEC4A | Additional_Macrophage | | | NM_194448.2 | 389-488 |
| CLK1 | nCounter_Kinase | | | NM_004071.3 | 759-858 |
| CNOT10 | Additional_House_Keeping_Genes | | | NM_001256741.1 | 1963-2062 |
| CNOT4 | Additional_House_Keeping_Genes | | | NM_001190848.1 | 796-895 |
| COG7 | Additional_House_Keeping_Genes | | | NM_153603.3 | 1493-1592 |
| COL8A2 | Additional_Macrophage | | | NM_005202.1 | 4001-4100 |
| CR2 | nCounter_Immunology | | | NM_001006658.1 | 486-585 |
| CREB3L2 | Additional_Lymph2Cx | | | NM_001253775.1 | 497-596 |
| CREBBP | Additional_Recurrently_mutated_in_DLBCL | | | NM_004380.2 | 1302-1401 |
| CSNK1D | nCounter_Kinase | | | NM_001893.3 | 1033-1132 |
| CTLA4 | nCounter_Immunology | RNAseq | | NM_005214.3 | 406-505 |
| CXCL13 | nCounter_Immunology | | | NM_006419.2 | 211-310 |
| CXCL5 | Additional_Macrophage | | | NM_002994.3 | 251-350 |
| CXCR3 | nCounter_Immunology | | | NM_001504.1 | 81-180 |
| CXCR5 | Additional_T_cell_subset | | | NM_001716.3 | 2619-2718 |
| CYB5R2 | Additional_Lymph2Cx | | | NM_016229.3 | 367-466 |
| CYP2U1 | RNAseq | | | NM_183075.2 | 2205-2304 |
| DAPK2 | nCounter_Kinase | | | NM_014326.3 | 754-853 |
| DCLK1 | nCounter_Kinase | | | NM_004734.4 | 1572-1671 |
| DDR2 | nCounter_Kinase | | | NM_006182.2 | 1707-1806 |
| DDX50 | Additional_House_Keeping_Genes | | | NM_024045.1 | 1186-1285 |
| DHX16 | Additional_House_Keeping_Genes | | | NM_001164239.1 | 2491-2590 |
| DLL4 | nCounter_PanCancer_Pathway | | | NM_019074.2 | 894-993 |
| DNAJC14 | Additional_House_Keeping_Genes | | | NM_032364.5 | 1167-1266 |
| DPY19L2P2 | RNAseq | | | NR_003561.2 | 3313-3412 |
| DSTYK | nCounter_Kinase | | | NM_015375.1 | 5206-5305 |
| DUSP4 | nCounter_Immunology | nCounter_PanCancer_Pathway | | NM_057158.2 | 3116-3215 |
| DYRK3 | nCounter_Kinase | | | NM_003582.2 | 1311-1410 |
| E2F1 | nCounter_PanCancer_Pathway | | | NM_005225.1 | 936-1035 |
| E2F2 | RNAseq | | | NM_004091.2 | 3606-3705 |
| EDC3 | Additional_House_Keeping_Genes | | | NM_001142443.1 | 1025-1124 |
| EEF1G | Additional_House_Keeping_Genes | | | NM_001404.4 | 1151-1250 |
| EGR2 | nCounter_Immunology | RNAseq | | NM_000399.3 | 1892-1991 |

| | | | | | |
|---|---|---|---|---|---|
| EIF2B4 | Additional_House_Keeping_Genes | | | NM_172195.3 | 1391-1490 |
| EIF3B | RNAseq | | | NM_001037283.1 | 961-1060 |
| ENDOG | nCounter_PanCancer_Pathway | | | NM_004435.2 | 695-794 |
| EP300 | Additional_Recurrently_mutated_in_DLBCL | | | NM_001429.2 | 716-815 |
| EPB41L4B | RNAseq | | | NM_018424.2 | 1536-1635 |
| EPHA1 | nCounter_Kinase | | | NM_005232.3 | 2076-2175 |
| EPHA3 | nCounter_Kinase | | | NM_005233.5 | 1280-1379 |
| EPHB6 | nCounter_Kinase | | | NM_004445.3 | 3785-3884 |
| ERCC3 | Additional_House_Keeping_Genes | | | NM_000122.1 | 1951-2050 |
| ERCC6 | nCounter_PanCancer_Pathway | | | NM_000124.2 | 3236-3335 |
| ETFA | RNAseq | | | NM_001127716.1 | 631-730 |
| EZH2 | Additional_Recurrently_mutated_in_DLBCL | | | NM_004456.3 | 191-290 |
| FAS | nCounter_Immunology | nCounter_PanCancer_Pathway | RNAseq | NM_000043.4 | 399-498 |
| FASLG | Additional_T_cell_subset | | | NM_000639.1 | 626-725 |
| FCER2 | Additional_Macrophage | | | NM_002002.4 | 421-520 |
| FCF1 | Additional_House_Keeping_Genes | | | NM_015962.4 | 229-328 |
| FER | nCounter_Kinase | | | NM_005246.1 | 611-710 |
| FGFR1 | nCounter_Kinase | nCounter_PanCancer_Pathway | | NM_015850.2 | 1336-1435 |
| FKBP4 | RNAseq | | | NM_002014.3 | 311-410 |
| FOXM1 | Additional_globally_associated_with_adverse_survival_PRECOG | | | NM_202002.1 | 1001-1100 |
| FOXO1 | Additional_T_cell_subset | | | NM_002015.3 | 1527-1626 |
| FOXO3 | Additional_T_cell_subset | | | NM_001455.2 | 1861-1960 |
| FOXP3 | nCounter_Immunology | | | NM_014009.3 | 1231-1330 |
| FOXRED2 | RNAseq | | | NM_001102371.1 | 2995-3094 |
| FYN | nCounter_Immunology | nCounter_Kinase | | NM_002037.3 | 766-865 |
| G6PD | Additional_House_Keeping_Genes | | | NM_000402.2 | 1156-1255 |
| GAPDH | Additional_House_Keeping_Genes | | | NM_002046.3 | 973-1072 |
| GAR1 | RNAseq | | | NM_032993.2 | 642-741 |
| GARS | RNAseq | | | NM_002047.2 | 1231-1330 |
| GNA13 | Additional_Recurrently_mutated_in_DLBCL | | | NM_006572.4 | 631-730 |
| GNAI2 | Additional_Recurrently_mutated_in_DLBCL | | | NM_002070.2 | 1001-1100 |
| GPATCH3 | Additional_House_Keeping_Genes | | | NM_022078.2 | 235-334 |
| GPR183 | nCounter_Immunology | | | NM_004951.3 | 406-505 |
| GRAMD1C | RNAseq | | | NM_017577.4 | 3385-3484 |
| GRK5 | nCounter_Kinase | | | NM_005308.2 | 1736-1835 |
| GSG2 | nCounter_Kinase | | | NM_031965.2 | 1836-1935 |
| GUSB | Additional_House_Keeping_Genes | | | NM_000181.1 | 1351-1450 |

| | | | | | |
|---|---|---|---|---|---|
| HDAC2 | nCounter_PanCancer_Pathway | | | NM_001527.1 | 931-1030 |
| HDAC3 | Additional_House_Keeping_Genes | | | NM_003883.3 | 353-452 |
| HIST1H3I | RNAseq | | | NM_003533.2 | 358-457 |
| HLA-DRB1 | nCounter_Immunology | | | NM_002124.3 | 748-847 |
| HPRT1 | Additional_House_Keeping_Genes | | | NM_000194.1 | 241-340 |
| HUNK | nCounter_Kinase | | | NM_014586.1 | 1281-1380 |
| ICA1 | RNAseq | | | NM_004968.3 | 1623-1722 |
| ICAM3 | nCounter_Immunology | | | NM_002162.3 | 1226-1325 |
| ICAM5 | nCounter_Immunology | | | NM_003259.3 | 328-427 |
| ICOS | nCounter_Immunology | RNAseq | | NM_012092.2 | 641-740 |
| ICOSLG | nCounter_Immunology | | | NM_015259.5 | 643-742 |
| IFI35 | nCounter_Immunology | | | NM_005533.3 | 416-515 |
| IFNAR2 | nCounter_Immunology | | | NM_000874.3 | 632-731 |
| IFNGR1 | nCounter_Immunology | | | NM_000416.1 | 1141-1240 |
| IFRD2 | RNAseq | | | NM_006764.3 | 1251-1350 |
| IKBKB | nCounter_Immunology | nCounter_Kinase | | NM_001556.1 | 1996-2095 |
| IKBKE | nCounter_Immunology | nCounter_Kinase | | NM_014002.2 | 2471-2570 |
| IKBKG | nCounter_Immunology | | | NM_003639.2 | 471-570 |
| IKZF2 | nCounter_Immunology | | | NM_001079526.1 | 3363-3462 |
| IL10 | Additional_Macrophage | | | NM_000572.2 | 231-330 |
| IL11RA | nCounter_Immunology | | | NM_147162.1 | 401-500 |
| IL12RB1 | nCounter_Immunology | | | NM_005535.1 | 449-548 |
| IL2 | nCounter_Immunology | | | NM_000586.2 | 301-400 |
| IL21 | nCounter_Immunology | RNAseq | | NM_021803.2 | 66-165 |
| IL21R | nCounter_Immunology | | | NM_021798.2 | 2081-2180 |
| IL27RA | Additional_Macrophage | | | NM_004843.3 | 1641-1740 |
| IL2RA | Additional_T_cell_subset | | | NM_000417.1 | 1001-1100 |
| IL2RB | nCounter_Immunology | | | NM_000878.2 | 1981-2080 |
| IL2RG | nCounter_Immunology | | | NM_000206.1 | 596-695 |
| IL32 | Additional_T_cell_subset | | | NM_004221.4 | 359-458 |
| IL6R | nCounter_Immunology | | | NM_000565.2 | 994-1093 |
| IL7R | nCounter_Immunology | | | NM_002185.3 | 401-500 |
| ILK | nCounter_Kinase | | | NM_004517.2 | 984-1083 |
| IRF4 | Additional_Lymph2Cx | | | NM_002460.1 | 326-425 |
| ISY1 | Additional_Lymph2Cx | | | NM_020701.2 | 87-186 |
| ITGA2B | Additional_Megakaryocyte | | | NM_000419.3 | 741-840 |
| ITGA4 | nCounter_Immunology | | | NM_000885.4 | 976-1075 |
| ITGAL | nCounter_Immunology | | | NM_002209.2 | 1114-1213 |
| ITGAX | nCounter_Immunology | | | NM_000887.3 | 701-800 |
| ITGB2 | nCounter_Immunology | | | NM_000211.2 | 521-620 |
| ITGB3 | Additional_Megakaryocyte | | | NM_000212.2 | 4486-4585 |

| | | | | | |
|---|---|---|---|---|---|
| ITK | nCounter_Kinase | | | NM_005546.3 | 3431-3530 |
| ITPKB | Additional_Lymph2Cx | | | NM_002221.3 | 79-178 |
| JAK1 | nCounter_Immunology | nCounter_Kinase | | NM_002227.1 | 286-385 |
| JAK2 | nCounter_Immunology | nCounter_Kinase | | NM_004972.2 | 456-555 |
| JAK3 | nCounter_Immunology | nCounter_Kinase | | NM_000215.2 | 1716-1815 |
| KLRB1 | nCounter_Immunology | | | NM_002258.2 | 86-185 |
| KMT2D | Additional_Recurrently_mutated_in_DLBCL | | | NM_003482.3 | 6071-6170 |
| LAMB3 | nCounter_PanCancer_Pathway | | | NM_000228.2 | 696-795 |
| LAMC2 | nCounter_PanCancer_Pathway | | | NM_005562.2 | 2820-2919 |
| LAMC3 | nCounter_PanCancer_Pathway | | | NM_006059.3 | 2091-2190 |
| LCP2 | nCounter_Immunology | | | NM_005565.3 | 1828-1927 |
| LIF | nCounter_Immunology | | | NM_002309.3 | 1241-1340 |
| LIFR | Additional_Others | | | NM_002310.3 | 2996-3095 |
| LIMD1 | Additional_Lymph2Cx | | | NM_014240.2 | 2926-3025 |
| LIMK1 | nCounter_Kinase | | | NM_002314.3 | 3134-3233 |
| lnc_AC012652 | Additional_lncRNA | | | lnc_AC012652.1 | 121-220 |
| lnc_ACPL2_3 | Additional_lncRNA | | | lnc_ACPL2_3.1 | 468-567 |
| lnc_ADRBK2_5 | Additional_lncRNA | | | lnc_ADRBK2_5.1 | 996-1095 |
| lnc_AL353597 | Additional_lncRNA | | | lnc_AL353597.1 | 249-348 |
| lnc_BCL11A_2 | Additional_lncRNA | | | lnc_BCL11A_2.1 | 740-839 |
| lnc_BCL2L11_3 | Additional_lncRNA | | | lnc_BCL2L11_3.1 | 269-368 |
| lnc_BCOR_5 | Additional_lncRNA | | | lnc_BCOR_5.1 | 157-256 |
| lnc_C2orf78_2 | Additional_lncRNA | | | lnc_C2orf78_2.1 | 219-318 |
| lnc_C5_1 | Additional_lncRNA | | | lnc_C5_1.1 | 479-578 |
| lnc_CD58_1 | Additional_lncRNA | | | lnc_CD58_1.1 | 578-677 |
| lnc_CD58_2 | Additional_lncRNA | | | lnc_CD58_2.1 | 201-300 |
| lnc_CLEC19A | Additional_lncRNA | | | lnc_CLEC19A.1 | 212-311 |
| lnc_CRHR1_4 | Additional_lncRNA | | | lnc_CRHR1_4.1 | 622-721 |
| lnc_DIP2A_1 | Additional_lncRNA | | | lnc_DIP2A_1.1 | 248-347 |
| lnc_DLG5_1 | Additional_lncRNA | | | lnc_DLG5_1.1 | 695-794 |
| lnc_DLG5_3 | Additional_lncRNA | | | lnc_DLG5_3.1 | 644-743 |
| lnc_EGR3_2 | Additional_lncRNA | | | lnc_EGR3_2.1 | 511-610 |
| lnc_EIF2S2_4 | Additional_lncRNA | | | lnc_EIF2S2_4.1 | 163-262 |
| lnc_FAS_1 | Additional_lncRNA | | | lnc_FAS_1.1 | 638-737 |
| lnc_FAS_2 | Additional_lncRNA | | | lnc_FAS_2.1 | 1506-1605 |
| lnc_FBXO43_8 | Additional_lncRNA | | | lnc_FBXO43_8.1 | 60-159 |
| lnc_GALK1_4 | Additional_lncRNA | | | lnc_GALK1_4.1 | 109-208 |
| lnc_GLUD1_8 | Additional_lncRNA | | | lnc_GLUD1_8.1 | 162-261 |
| lnc_GMIP_1 | Additional_lncRNA | | | lnc_GMIP_1.1 | 237-336 |
| lnc_HYAL4_4 | Additional_lncRNA | | | lnc_HYAL4_4.1 | 567-666 |
| lnc_KIAA0020_9 | Additional_lncRNA | | | lnc_KIAA0020_9.1 | 891-990 |

| | | | | | |
|---|---|---|---|---|---|
| lnc_LRRC3_6 | Additional_lncRNA | | | lnc_LRRC3_6.1 | 265-364 |
| lnc_LY75_1 | Additional_lncRNA | | | lnc_LY75_1.1 | 747-846 |
| lnc_MYC_1 | Additional_lncRNA | | | lnc_MYC_1.1 | 2899-2998 |
| lnc_NRXN3_4 | Additional_lncRNA | | | lnc_NRXN3_4.1 | 558-657 |
| lnc_NT5C_1 | Additional_lncRNA | | | lnc_NT5C_1.1 | 133-232 |
| lnc_PARD3B_1 | Additional_lncRNA | | | lnc_PARD3B_1.1 | 8263-8362 |
| lnc_PBX4_1 | Additional_lncRNA | | | lnc_PBX4_1.1 | 410-509 |
| lnc_PDIA4_3 | Additional_lncRNA | | | lnc_PDIA4_3.1 | 247-346 |
| lnc_PDYN_1 | Additional_lncRNA | | | lnc_PDYN_1.1 | 590-689 |
| lnc_PHF1_1 | Additional_lncRNA | | | lnc_PHF1_1.1 | 287-386 |
| lnc_PTTG1_1 | Additional_lncRNA | | | lnc_PTTG1_1.1 | 690-789 |
| lnc_PTTG1_9 | Additional_lncRNA | | | lnc_PTTG1_9.1 | 774-873 |
| lnc_PYGO2_2 | Additional_lncRNA | | | lnc_PYGO2_2.1 | 1187-1286 |
| lnc_RAPH1_2 | Additional_lncRNA | | | lnc_RAPH1_2.1 | 1777-1876 |
| lnc_RP11_478C19 | Additional_lncRNA | | | lnc_RP11_478C19.1 | 99-198 |
| lnc_RPS6KL1_1 | Additional_lncRNA | | | lnc_RPS6KL1_1.1 | 951-1050 |
| lnc_SEMA4D_5 | Additional_lncRNA | | | lnc_SEMA4D_5.1 | 163-262 |
| lnc_SERINC1_3 | Additional_lncRNA | | | lnc_SERINC1_3.1 | 951-1050 |
| lnc_SNX11_7 | Additional_lncRNA | | | lnc_SNX11_7.1 | 653-752 |
| lnc_SPACA5_1 | Additional_lncRNA | | | lnc_SPACA5_1.1 | 40-139 |
| lnc_STAMBPL1_1 | Additional_lncRNA | | | lnc_STAMBPL1_1.1 | 412-511 |
| lnc_STC1_2 | Additional_lncRNA | | | lnc_STC1_2.1 | 371-470 |
| lnc_TDRD7_1 | Additional_lncRNA | | | lnc_TDRD7_1.1 | 1509-1608 |
| lnc_TMEM194B_5 | Additional_lncRNA | | | lnc_TMEM194B_5.1 | 1162-1261 |
| lnc_TMEM63A_1 | Additional_lncRNA | | | lnc_TMEM63A_1.1 | 87-186 |
| lnc_TPCN2_5 | Additional_lncRNA | | | lnc_TPCN2_5.1 | 138-237 |
| lnc_TRA2A_5 | Additional_lncRNA | | | lnc_TRA2A_5.1 | 349-448 |
| lnc_ZFP36L2_2 | Additional_lncRNA | | | lnc_ZFP36L2_2.1 | 184-283 |
| lnc_ZNF143_3 | Additional_lncRNA | | | lnc_ZNF143_3.1 | 207-306 |
| lnc_ZNF385C_6 | Additional_lncRNA | | | lnc_ZNF385C_6.1 | 803-902 |
| LRRC2 | RNAseq | | | NM_024512.4 | 861-960 |
| LYAR | RNAseq | | | NM_001145725.1 | 231-330 |
| MAD2L2 | nCounter_PanCancer_Pathway | | | NM_001127325.1 | 291-390 |
| MAF | nCounter_Immunology | | | NM_005360.4 | 889-988 |
| MALT1 | nCounter_Immunology | | | NM_006785.2 | 910-1009 |
| MAML3 | Additional_Lymph2Cx | | | NM_018717.4 | 1351-1450 |
| MAP2K4 | nCounter_Kinase | | | NM_003010.2 | 772-871 |
| MAP3K8 | nCounter_Kinase | | | NM_005204.2 | 2051-2150 |
| MAP4K4 | nCounter_Immunology | | | NM_004834.3 | 3316-3415 |

| | | | | | |
|---|---|---|---|---|---|
| MAPK12 | nCounter_Kinase | nCounter_PanCancer_Pathway | | NM_002969.3 | 426-525 |
| MAPK3 | nCounter_Kinase | | | NM_001040056.1 | 581-680 |
| MARCO | Additional_Macrophage | | | NM_006770.3 | 1435-1534 |
| MARK1 | nCounter_Kinase | | | NM_018650.2 | 1871-1970 |
| MAST3 | nCounter_Kinase | | | NM_015016.1 | 4061-4160 |
| MCL1 | nCounter_Immunology | | | NM_021960.3 | 1261-1360 |
| MCM4 | nCounter_PanCancer_Pathway | | | NM_182746.1 | 1201-1300 |
| MCM7 | nCounter_PanCancer_Pathway | | | NM_182776.1 | 1326-1425 |
| MEF2B | Additional_Recurrently_mutated_in_DLBCL | | | NM_001145785.1 | 10-109 |
| MEIS2 | RNAseq | | | NM_002399.2 | 961-1060 |
| MFNG | nCounter_PanCancer_Pathway | | | NM_002405.2 | 1682-1781 |
| MME | Additional_Lymph2Cx | | | NM_000902.2 | 5060-5159 |
| CD206 | Additional_Macrophage | | | NM_002438.2 | 526-625 |
| MRPL46 | RNAseq | | | NM_022163.3 | 301-400 |
| MRPS5 | Additional_House_Keeping_Genes | | | NM_031902.3 | 391-490 |
| MRTO4 | RNAseq | | | NM_016183.3 | 309-408 |
| MS4A1 | nCounter_Immunology | | | NM_152866.2 | 621-720 |
| MST1R | nCounter_Kinase | | | NM_002447.1 | 3301-3400 |
| MTMR14 | Additional_House_Keeping_Genes | | | NM_022485.3 | 721-820 |
| MTOR | Additional_BCR_signal | | | NM_004958.3 | 1866-1965 |
| MUSK | nCounter_Kinase | | | NM_005592.1 | 1746-1845 |
| MYB | Additional_T_cell_subset | | | NM_001130173.1 | 184-283 |
| MYBBP1A | RNAseq | | | NM_001105538.1 | 3896-3995 |
| MYBL1 | Additional_Lymph2Cx | | | NM_001080416.3 | 1031-1130 |
| MYC | nCounter_PanCancer_Pathway | RNAseq | | NM_002467.3 | 1611-1710 |
| MYCN | nCounter_PanCancer_Pathway | | | NM_005378.4 | 1546-1645 |
| MYD88 | Additional_Recurrently_mutated_in_DLBCL | | | NM_002468.3 | 2146-2245 |
| C19orf10 | RNAseq | | | NM_019107.3 | 650-749 |
| NEK6 | nCounter_Kinase | | | NM_014397.5 | 659-758 |
| NFATC2 | nCounter_Immunology | | | NM_012340.3 | 1816-1915 |
| NFATC3 | nCounter_Immunology | | | NM_004555.2 | 2191-2290 |
| NFKB1 | Additional_BCR_signal | | | NM_003998.2 | 1676-1775 |
| NFKB2 | nCounter_Immunology | | | NM_001077493.1 | 1062-1161 |
| NFKBIA | nCounter_Immunology | | | NM_020529.1 | 946-1045 |
| NOD2 | nCounter_Immunology | | | NM_022162.2 | 559-658 |
| NOL7 | Additional_House_Keeping_Genes | | | NM_016167.3 | 336-435 |
| NOP14 | RNAseq | | | XM_005248037.1 | 2326-2425 |
| NOTCH1 | nCounter_Immunology | | | NM_017617.3 | 8212-8311 |
| NPR2 | nCounter_Kinase | | | NM_003995.3 | 3268-3367 |
| NRBP1 | nCounter_Kinase | | | NM_013392.2 | 163-262 |

| | | | | | |
|---|---|---|---|---|---|
| NUBP1 | Additional_House_Keeping_Genes | | | NM_001278506.1 | 305-404 |
| NUDT4 | RNAseq | | | NM_199040.2 | 4108-4207 |
| OAZ1 | Additional_House_Keeping_Genes | | | NM_004152.2 | 314-413 |
| OBSCN | nCounter_Kinase | | | NM_001098623.2 | 1309-1408 |
| PAICS | RNAseq | | | NM_006452.3 | 2948-3047 |
| PAX5 | nCounter_Immunology | | | NM_016734.1 | 2289-2388 |
| PBK | nCounter_Kinase | | | NM_018492.2 | 1588-1687 |
| PDCD1 | nCounter_Immunology | | | NM_005018.1 | 176-275 |
| PDE4D | RNAseq | | | NM_006203.4 | 5581-5680 |
| PDK3 | nCounter_Kinase | | | NM_005391.1 | 586-685 |
| PDXP | RNAseq | | | NM_020315.4 | 1399-1498 |
| PF4 | Additional_Megakaryocyte | | | NM_002619.2 | 1-100 |
| PIK3CA | Additional_BCR_signal | | | NM_006218.2 | 2446-2545 |
| PIK3CB | Additional_BCR_signal | | | NM_006219.1 | 2946-3045 |
| PIK3CD | Additional_BCR_signal | | | NM_005026.3 | 2979-3078 |
| PIK3CG | Additional_BCR_signal | | | NM_002649.2 | 2126-2225 |
| PIM2 | Additional_Lymph2Cx | | | NM_006875.2 | 621-720 |
| PLAU | nCounter_Immunology | | | NM_002658.2 | 794-893 |
| POLR1B | nCounter_Immunology | | | NM_019014.3 | 3321-3420 |
| POLR2A | Additional_House_Keeping_Genes | | | NM_000937.2 | 3776-3875 |
| POLR2H | nCounter_PanCancer_Pathway | | | NM_001278698.1 | 941-1040 |
| PPBP | Additional_Macrophage | | | NM_002704.2 | 331-430 |
| PPIA | Additional_House_Keeping_Genes | | | NM_021130.3 | 201-300 |
| PRDM1 | Additional_Recurrently_mutated_in_DL BCL | | | NM_182907.1 | 311-410 |
| PRKACB | nCounter_Kinase | | | NM_182948.2 | 806-905 |
| PRKCB | Additional_BCR_signal | | | NM_212535.1 | 1751-1850 |
| PRKCH | nCounter_Kinase | | | NM_006255.3 | 851-950 |
| PRKD1 | nCounter_Kinase | | | NM_002742.1 | 1491-1590 |
| PRKD2 | nCounter_Kinase | | | NM_001079882.1 | 1651-1750 |
| PRPF38A | Additional_House_Keeping_Genes | | | NM_032864.3 | 336-435 |
| PSAT1 | RNAseq | | | NM_021154.3 | 1446-1545 |
| PSPH | RNAseq | | | XM_005271773.1 | 803-902 |
| PTK2B | nCounter_Kinase | | | NM_004103.3 | 736-835 |
| PTPN6 | nCounter_Immunology | | | NM_002831.5 | 1735-1834 |
| PTPRC_all | nCounter_Immunology | | | NM_080923.2 | 155-254 |
| R3HDM1 | Additional_Lymph2Cx | | | NM_015361.2 | 1276-1375 |
| RAB27A | RNAseq | | | NM_183236.1 | 1581-1680 |
| RAB7L1 | Additional_Lymph2Cx | | | NM_001135664.1 | 786-885 |
| RABL2A | RNAseq | | | NM_007082.3 | 316-415 |
| RELA | nCounter_Immunology | | | NM_021975.2 | 361-460 |
| RELB | nCounter_Immunology | | | NM_006509.3 | 961-1060 |

| | | | | | |
|---|---|---|---|---|---|
| ROCK2 | nCounter_Kinase | | | NM_004850.3 | 3141-3240 |
| ROR2 | nCounter_Kinase | | | NM_004560.3 | 3383-3482 |
| RPL19 | Additional_House_Keeping_Genes | | | NM_000981.3 | 316-415 |
| RPS6KA3 | nCounter_Kinase | | | NM_004586.2 | 278-377 |
| RPS6KB1 | Additional_BCR_signal | | | NM_001272042.1 | 661-760 |
| S100A9 | nCounter_Immunology | | | NM_002965.2 | 76-175 |
| S1PR2 | Additional_Lymph2Cx | | | NM_004230.2 | 186-285 |
| SAP130 | Additional_House_Keeping_Genes | | | NM_024545.3 | 3091-3190 |
| SARDH | RNAseq | | | NM_001134707.1 | 805-904 |
| SCYL2 | nCounter_Kinase | | | NM_017988.4 | 2416-2515 |
| SCYL3 | nCounter_Kinase | | | NM_020423.4 | 611-710 |
| SDHA | Additional_House_Keeping_Genes | | | NM_004168.1 | 231-330 |
| SERPINA9 | Additional_Lymph2Cx | | | NM_001042518.1 | 1156-1255 |
| SF3A3 | Additional_House_Keeping_Genes | | | NM_006802.2 | 2061-2160 |
| SGK1 | nCounter_Kinase | | | NM_005627.2 | 1791-1890 |
| SH2D1A | nCounter_Immunology | | | NM_002351.4 | 496-595 |
| SHMT2 | RNAseq | | | NM_001166356.1 | 1461-1560 |
| SLAMF1 | nCounter_Immunology | | | NM_003037.2 | 581-680 |
| SLAMF6 | nCounter_Immunology | | | NM_001184714.1 | 1033-1132 |
| SLC2A1 | nCounter_Immunology | | | NM_006516.2 | 2501-2600 |
| SMAD2 | Additional_T_cell_subset | | | NM_005901.5 | 1679-1778 |
| SNUPN | RNAseq | | | NM_001042581.1 | 885-984 |
| SOCS1 | nCounter_Immunology | | | NM_003745.1 | 1026-1125 |
| SPI1 | Additional_T_cell_subset | | | NM_003120.1 | 731-830 |
| SRM | RNAseq | | | NM_003132.2 | 255-354 |
| STAT2 | nCounter_Immunology | | | NM_005419.3 | 1391-1490 |
| STAT4 | nCounter_Immunology | | | NM_003151.2 | 790-889 |
| STAT5B | nCounter_Immunology | | | NM_012448.3 | 1411-1510 |
| STK17A | nCounter_Kinase | | | NM_004760.1 | 1106-1205 |
| STK17B | nCounter_Kinase | | | NM_004226.3 | 822-921 |
| STK24 | nCounter_Kinase | | | NM_001032296.1 | 1206-1305 |
| STK32B | nCounter_Kinase | | | NM_018401.1 | 926-1025 |
| SULF2 | RNAseq | | | NM_001161841.1 | 1207-1306 |
| SUMO4 | RNAseq | | | NM_001002255.1 | 157-256 |
| SYK | Additional_BCR_signal | | | NM_003177.5 | 809-908 |
| TAB2 | RNAseq | | | NM_015093.3 | 1378-1477 |
| TAF1 | nCounter_Kinase | | | NM_004606.4 | 1067-1166 |
| TBC1D4 | RNAseq | | | NM_014832.2 | 3131-3230 |
| TBP | Additional_House_Keeping_Genes | | | NM_001172085.1 | 588-687 |
| TBX21 | nCounter_Immunology | | | NM_013351.1 | 891-990 |
| TCF7 | nCounter_Immunology | | | NM_201633.2 | 520-619 |

| | | | | | |
|---|---|---|---|---|---|
| TERT | RNAseq | | | NM_198253.1 | 2571-2670 |
| TFAP4 | RNAseq | | | XM_011522634.1 | 121-220 |
| TGFB | Additional_Macrophage | | | NM_000660.5 | 1846-1945 |
| TGFBR1 | nCounter_Immunology | nCounter_Kinase | | NM_004612.2 | 4281-4380 |
| TIGIT | nCounter_Immunology | | | NM_173799.2 | 1969-2068 |
| TIMM8B | RNAseq | | | NR_028383.1 | 511-610 |
| TLK1 | nCounter_Kinase | | | NM_001136554.1 | 509-608 |
| TLK2 | Additional_House_Keeping_Genes | | | XM_011524223.1 | 384-483 |
| TMEM173 | nCounter_Immunology | | | NM_198282.1 | 726-825 |
| TMUB2 | Additional_House_Keeping_Genes | | | NM_024107.2 | 1486-1585 |
| TNFAIP3 | Additional_Recurrently_mutated_in_DLBCL | | | NM_006290.2 | 261-360 |
| TNFRSF13B | Additional_Lymph2Cx | | | NM_012452.2 | 161-260 |
| TNFRSF14 | Additional_Recurrently_mutated_in_DLBCL | | | NM_003820.3 | 660-759 |
| TNFRSF18 | Additional_T_cell_subset | | | NM_148901.1 | 303-402 |
| TNFRSF4 | nCounter_Immunology | | | NM_003327.3 | 446-545 |
| TNFRSF9 | nCounter_Immunology | | | NM_001561.4 | 256-355 |
| TNIK | nCounter_Kinase | | | NM_001161560.1 | 246-345 |
| TP53 | Additional_Recurrently_mutated_in_DLBCL | | | NM_000546.2 | 1331-1430 |
| TRAF1 | nCounter_Immunology | RNAseq | | NM_005658.3 | 3736-3835 |
| TRAF3 | nCounter_Immunology | | | NM_145725.2 | 721-820 |
| TRAF5 | nCounter_Immunology | | | NM_001033910.2 | 1691-1790 |
| TRIM39 | Additional_House_Keeping_Genes | | | NM_021253.3 | 3141-3240 |
| TRIM56 | Additional_Lymph2Cx | | | NM_030961.1 | 2571-2670 |
| TRIM65 | RNAseq | | | NM_173547.2 | 986-1085 |
| TRIO | nCounter_Kinase | | | NM_007118.2 | 1736-1835 |
| TTC31 | nCounter_PanCancer_Pathway | | | NR_027749.1 | 411-510 |
| TUBB | Additional_House_Keeping_Genes | | | NM_178014.2 | 1956-2055 |
| TXK | nCounter_Kinase | | | NM_003328.1 | 801-900 |
| UBXN4 | Additional_Lymph2Cx | | | NM_014607.3 | 344-443 |
| USP39 | Additional_House_Keeping_Genes | | | NM_001256725.1 | 807-906 |
| VCAM1 | nCounter_Immunology | | | NM_001078.3 | 2536-2635 |
| WDR55 | Additional_Lymph2Cx | | | NM_017706.4 | 816-915 |
| XLOC_001814 | Additional_lncRNA | | | XLOC_001814.1 | 444-543 |
| XLOC_002470 | Additional_lncRNA | | | XLOC_002470.1 | 147-246 |
| XLOC_006922 | Additional_lncRNA | | | XLOC_006922.1 | 316-415 |
| XLOC_006923 | Additional_lncRNA | | | XLOC_006923.1 | 1355-1454 |
| XLOC_006924 | Additional_lncRNA | | | XLOC_006924.1 | 636-735 |
| XLOC_006925 | Additional_lncRNA | | | XLOC_006925.1 | 192-291 |
| XLOC_007217 | Additional_lncRNA | | | XLOC_007217.1 | 829-928 |

| | | | | | |
|---|---|---|---|---|---|
| XLOC_007218 | Additional_lncRNA | | | XLOC_007218.1 | 592-691 |
| XLOC_007219 | Additional_lncRNA | | | XLOC_007219.1 | 104-203 |
| XYLB | RNAseq | | | NM_005108.3 | 1516-1615 |
| YES1 | nCounter_Kinase | | | NM_005433.3 | 266-365 |
| ZAP70 | nCounter_Immunology | nCounter_Kinase | | NM_001079.3 | 1176-1275 |
| ZC3H14 | Additional_House_Keeping_Genes | | | NM_207662.3 | 1207-1306 |
| ZEB1 | nCounter_Immunology | | | NM_001128128.1 | 1451-1550 |
| ZKSCAN5 | Additional_House_Keeping_Genes | | | NM_014569.3 | 3689-3788 |
| ZNF143 | Additional_House_Keeping_Genes | | | NM_003442.5 | 926-1025 |
| ZNF346 | Additional_House_Keeping_Genes | | | NM_012279.2 | 761-860 |

### Table S8. Multivariate analysis in the validation cohort

| Category | Group | Hazard ratio | 95%CI | p.value |
|---|---|---|---|---|
| DMS score | Low (vs. High) | 2.76 | (1.11-6.82) | 2.83E-02 |
| IPI | High (vs. Low) | 5.17 | (2.13-12.56) | 2.83E-04 |
| Lymph2Cx | ABC (vs. GCB) | 1.44 | (0.57-3.65) | 4.38E-01 |
| | Unclassified (vs. GCB) | 1.46 | (0.42-5.02) | 5.50E-01 |

### Table S9. Multivariate analysis in Schmitz et al. NEJM 2018

| Category | Group | Hazard ratio | 95%CI | p.value |
|---|---|---|---|---|
| Genetic Subtype | BN2 (vs. EZB) | 1.25 | (0.65-2.41) | 5.03.E-01 |
| | MCD (vs. EZB) | 2.77 | (1.42-5.38) | 2.70.E-03 |
| | N1 (vs. EZB) | 3.41 | (1.27-9.15) | 1.48.E-02 |
| | Others (vs. EZB) | 1.00 | (0.60-1.66) | 9.90.E-01 |
| DMS score | Low (vs. High) | 2.15 | (1.43-3.23) | 2.18.E-04 |

### Table S10. Multivariate analysis in Reddy et al. Cell 2017

| Category | Group | Hazard ratio | 95%CI | p.value |
|---|---|---|---|---|
| COO classification | ABC (vs. GCB) | 0.97 | (0.70-1.34) | 8.39.E-01 |
| | Unclassified (vs. GCB) | 1.18 | (0.75-1.87) | 4.79.E-01 |
| Genomic risk model | High risk (vs. Low risk) | 7.70 | (4.37-13.6) | 1.54.E-12 |
| | Medium risk (vs. Low risk) | 4.13 | (2.27-7.54) | 3.74.E-06 |
| DMS score | Low (vs. High) | 1.85 | (1.35-2.53) | 1.26.E-04 |
